# Translational regulation of GAD1 identified by circadian and light-responsive ribosome-bound transcriptome analysis in the mouse hypothalamic suprachiasmatic nucleus

**DOI:** 10.64898/2026.01.26.701657

**Authors:** Xinyan Shao, Sui-Wen Hsiao, Chenyang Xiao, Huihua Zhou, Youichi Tanaka, Tom Macpherson, Yuichi Shichino, Shintaro Iwasaki, Emi Hasegawa, Masao Doi

## Abstract

The suprachiasmatic nucleus (SCN) serves as the master circadian pacemaker, integrating regulation of clock genes in the SCN has been extensively studied, whether gene expression in the SCN is regulated at the level of mRNA translation has remained largely unexplored. Here, we report the first ribosome-profiling (Ribo-seq) dataset generated from the mouse SCN, enabling a genome-wide assessment of translational regulation in this central circadian clock. Combined time-series Ribo-seq and RNA-seq analyses identified 385 genes that exhibit rhythmic ribosome binding without corresponding oscillations in mRNA abundance, revealing widespread translational regulation in the SCN. Among these, light stimulation induced an approximately twofold increase in ribosome binding of *Gad1*, which encodes the major GABA-synthetic enzyme Gad67, despite only marginal changes in *Gad1* mRNA levels. Importantly, immunohistochemical analyses demonstrated light-dependent accumulation of Gad67 protein selectively in light-activated SCN neurons, establishing that translational regulation of *Gad1* gives rise to overt protein-level changes. In contrast, canonical clock genes were regulated predominantly at the transcriptional level and showed little evidence of translational modulation. Our data allow a systematic comparison between transcriptionally and translationally regulated genes, including their relative rhythm-amplitudes and phases, and thereby revealed translational control as a distinct modulatory layer shaping time-of-day-dependent and light-dependent gene expression in the SCN.

## INTRODUCTION

The suprachiasmatic nucleus (SCN) functions as the master pacemaker governing circadian behavior and physiology^1^. Although functional circadian clocks are present in many peripheral tissues, including the liver, kidney, and heart, the SCN occupies a unique position within the circadian system. Unlike other organs and cell types, SCN neurons receive direct photic input from the retina, enabling precise entrainment to the external light–dark cycle^2,3^. In addition, SCN neurons generate robust daily rhythms in neuronal activity that, through multiple direct and indirect output pathways, synchronize clocks across the body^4^. Consequently, the SCN resides at the apex of the circadian hierarchy, integrating environmental light information and coordinating peripheral clocks systemically. This distinctive role of the SCN as the central circadian clock is thought to arise from a combination of known and yet-to-be-elucidated mechanisms that extend beyond the core molecular clockwork shared by most cells^2,3^.

The γ-aminobutyric acid (GABA) is the major inhibitory neurotransmitter in the brain. GABA is primarily synthesized from glutamate by glutamate decarboxylase 67 kDa isoform (Gad67; encoded by *Gad1*) and/or glutamate decarboxylase 65 kDa isoform (Gad65; encoded by *Gad2*). Gad67 mediates synthesis of over 90% of basal GABA levels in the brain^5,6^. Of interest, nearly all SCN neurons are GABAergic^7^; however, temporal regulation of GABA availability by de novo Gad67/Gad65-dependent biosynthesis has received minor attention as a factor for quick regulation. A more common view is that GABA is loaded to synaptic vesicles, and vesicles are pooled inside presynaptic terminals before released into synaptic cleft (upon depolarization) by exocytosis^8^, so that nascent GABA-production is regarded as the prerequisite for constitutive GABA availability— not as the regulatory site for short-term regulation of signaling. On the other hand, the importance of the quantity control of Gad67 and Gad65 protein expression has been previously noted during development^9,10^ as well as in longer-term homeostatic regulation of GABA signaling^11^. It is interesting to note that, in the SCN, mRNA levels of *Gad1* and *Gad2* are likely under circadian regulation. Publicly available RNA-seq data^12^ report the rhythmicity fo *Gad2* but not *Gad1*. Previous microarray data^13^, on the other hand, report the opposite pattern; rhythmicity of *Gad1* but not *Gad2*. Previous *in situ* hybridization data^14^ identified low-amplitude day-night variations in *Gad2* mRNA expression but no significant difference in *Gad1* expression. Thus, although controversial, previous studies suggest that *Gad1*/*Gad2* expression in the SCN may be regulated on an hourly scale. Currently, no studies have characterized post-transcriptional regulation of *Gad1*/*Gad2* expression in the SCN. A potential response of *Gad1*/*Gad2* to light is also unstudied.

Ribo-seq is a powerful technique for studying in vivo translation at a genome-wide scale^15,16^. Time-series Ribo-seq analysis has been applied to illustrate circadian translatome in the liver^17,18^ and kidney^19^, two functionally different peripheral clock tissues. The studies in these organs verified that a number of transcripts exhibit rhythmic translation as a result of their rhythmic mRNA expression. Moreover, these studies revealed the existence of translationally rhythmic genes without concomitant mRNA abundance alteration. For example, in the liver, a specific subset of genes associated with iron metabolism are rhythmically translated from non-oscillating transcripts^16^; furthermore, cross-organ comparison between the liver and kidney shows that translationally only rhythmic genes are highly tissue-specific^19^. These observations in the peripheral tissue clocks raise interest in posing a parallel question on the translatome within the SCN. In particular, how light—an environmental cue uniquely processed by the SCN but not by the peripheral tissue clocks—impacts the SCN translatome remains to be explored.

In the present study, we report the first set of Ribo-seq data using the SCN as a source tissue. Anatomically, the SCN is a pair of nuclei situated near the third ventricle just above the optic chiasm, composed of only about 20,000 neurons. Like other nuclei within the hypothalamus, the anatomically small size of the SCN has been a constraint for material (tissue)-consuming study. In our study, we prepared microdissected SCN punches from six independent animals and pooled them as a single replicate. By performing time-series and light-illumination study, we found that *Gad1* expression is translationally regulated in the SCN. Immunohistological confirmation provided direct evidence for the protein-level increase of *Gad1*, or Gad67 after light stimulation in a specific set of light-activated SCN neurons. These data suggest a previously undescribed aspect of GABA availability regulation and expand our knowledge on the mode of gene expression regulation in the SCN. We also report Ribo-seq data for circadian clock genes in the SCN. Importantly, the data allow a systematic comparison between transcriptionally and translationally regulated genes in the SCN, including their relative oscillation amplitudes and temporal characteristics.

## RESULTS

### Time-series ribosome profiling in the SCN

In an attempt to seek for a potential post-transcriptional regulation of circadian gene expression, we asked if there are specific genes that oscillate in ribosome binding despite lacking mRNA abundance rhythm in the SCN. To test this hypothesis, time-series Ribo-seq was performed on the mouse SCN transcriptome across six time-points during a regular 12-h light/12-h dark cycle, in conjunction with RNA-seq examination (Fig. 1). Genes were considered rhythmic in RNA-seq or Ribo-seq if their permutation-based, adjusted p-value was < 0.05 (JTK_cycle^20^; see also Methods for details). Of the total 11,172 detected SCN genes, 502 genes were categorized as rhythmic in Ribo-seq (i.e., ribosome binding) and 428 in RNA-seq (i.e., mRNA abundance) (Fig. 1A, Venn diagram). 63 overlapping genes exhibited oscillation in both datasets, which include, as expected, clock genes such as *Per2*, *Cry1*, and *Nr1d1*. By simple assignment, 439 genes (502 minus 63) can be considered as “mRNA flat–ribosome binding rhythmic” genes.

**Figure 1.**
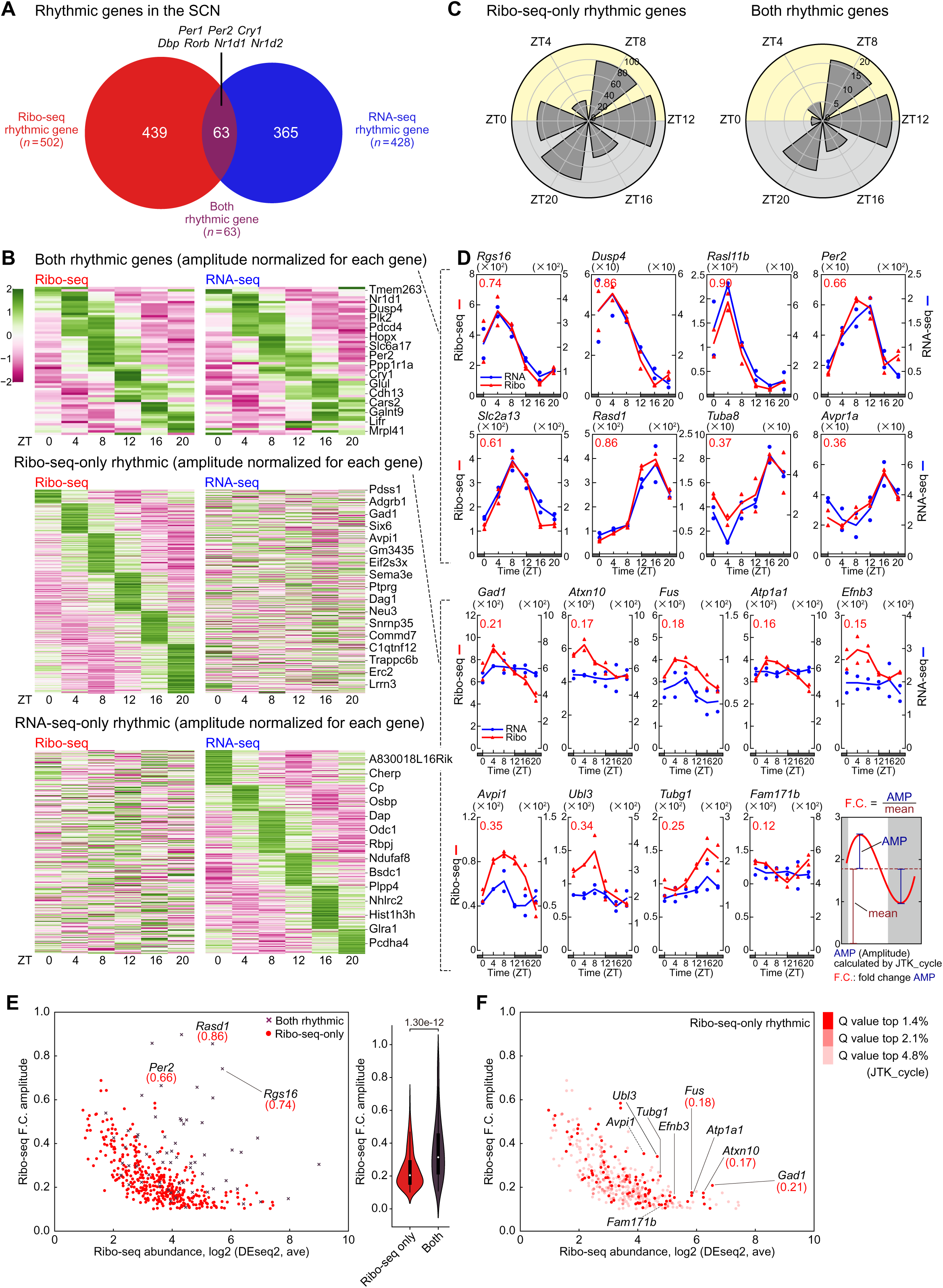
Identification of transcripts with circadian ribosome binding in the SCN. (**A**) Venn diagram showing the number of oscillating genes detected in RNA-seq (blue) and Ribo-seq (red) datasets selected by JTK_cycle (p < 0.05). Out of 11,172 genes, 428 are rhythmic in RNA-seq, 502 in Ribo-seq, and 63 in both. (**B**) Heatmaps of ribosome binding (Ribo-seq; left) and mRNA abundance (RNA-seq; right) rhythms for genes rhythmic in both (top), Ribo-seq specific (middle), and RNA-seq specific (bottom). Transcripts are sorted by phase. Amplitudes were standardized within each gene using Z-scores. RNA-seq-only rhythmic genes were selected as p > 0.2 for Ribo-seq; Ribo-seq-only were selected as p > 0.2 for RNA-seq. (**C**) Polar histogram plots of the peak phase for Ribo-seq specific and both rhythmic genes. (**D**) Representative oscillation profiles of both rhythmic genes and Ribo-seq-only rhythmic genes. Ribo-seq-only genes were selected based on their lack of evident RNA-seq rhythmicity. The red values on the graphs represent the F.C. amplitude of the Ribo-seq rhythm, calculated as the ratio of the JTK_cycle determined amplitude (AMP) to mean Ribo-seq abundance. (**E**) Scatter plots of both rhythmic and Ribo-seq-only rhythmic genes, plotted with the daily average Ribo-seq abundance on the *x*-axis and the Ribo-seq F.C. amplitude on the *y*-axis. Violin plots show the range of amplitude for each category. (**F**) Representative Ribo-seq-only rhythmic genes. The plots are the same as (E), with colors varying by q-value (JTK_cycle). The highlighted genes are plotted in (D). Statistics in (E), two-tailed unpaired *t*-test.

However, visual inspection of the corresponding Ribo-seq and RNA-seq profiles revealed that, in many cases, the apparent lack of rhythmicity at the mRNA level reflected borderline non-rhythmic assignments, with expression patterns still exhibiting oscillatory features (the borderline examples are shown in Supplementary Fig. 1). Within the Ribo-seq rhythmical genes, we therefore selectively isolated “Ribo-seq-only” rhythmic genes by choosing those which display rhythmicity (p < 0.05, JTK_cycle, as described) in Ribo-seq but clearly lack RNA-seq circadian variation (p > 0.2, JTK_cycle). This led us to identify 385 SCN genes (Fig. 1B, heatmap) that are statistically rhythmic in Ribo-seq without clear circadian variation in mRNA abundance.

In our heatmaps (Fig. 1B), transcripts are sorted according to phase, and oscillation amplitudes were normalized within individual genes for better visualization of their rhythmicity. In light of phase distribution (see also Fig. 1C, circular histogram), Ribo-seq-only rhythmic genes exhibit peak expression mainly either in the late phase of the light period, Zeitgeber time (ZT)8–12, or in the late night period, ZT20–0 (by definition, ZT0 and ZT12 refer to light on and off times, respectively). Gene transcripts that are rhythmic in both Ribo-seq and RNA-seq (hereafter referred to as “both rhythmic genes”) exhibit phase distributions more enriched at ZT8–12 (Fig. 1C), implying differences in phase control.

Additionally, we observed that 317 genes can be categorized as RNA-seq-only rhythmic genes (p > 0.2 in Ribo-seq, p < 0.05 in RNA-seq, JTK_cycle), indicating that a significant portion of rhythmic genes identified by RNA-seq (317 out of 365) do not exhibit corresponding rhythms in ribosome-bound transcripts. These genes are broadly distributed around the clock in respect to RNA-seq phase (Fig. 1B; see also Supplementary Fig. 2A for circular histogram).

### Oscillation amplitudes of ribosome-bound transcripts

Next, we turned our attention to the amplitude of Ribo-seq rhythmicity (Fig. 1D). To quantitatively compare the amplitude magnitude, we normalized the JTK_cycle-determined amplitude (AMP) to the mean daily abundance for each gene. Namely, we calculated the ratio (or fold change, F.C.) of AMP to mean Ribo-seq expression (see schematic in the bottom-right panel in Fig. 1D). Interestingly, we found that Ribo-seq-only rhythmic genes exhibited weaker amplitudes compared to the more robust rhythms of both rhythmic genes. For example, the magnitudes of the normalized F.C. amplitudes of the Ribo-seq-only rhythmic genes— *Gad1*, *Atxn10*, *Atp1a1*, *Fus, Efnb3*, *Avpi1*, *Ubl3*, *Tubg1*, and *Fam171b* —were 0.21, 0.17, 0.16, 0.18, 0.15, 0.35, 0.34, 0.25, and 0.12, respectively. On the other hand, the corresponding values for the both rhythmic genes — *Per2*, *Rgs16*, *Dusp4*, *Rasl11b*, *Slc2a13*, *Rasd1*, *Tuba8*, and *Avpr1a*— were 0.66, 0.74, 0.86, 0.99, 0.61, 0.86, 0.37, and 0.36, respectively (Fig. 1D, compare the red values on the graphs for each respective gene). This trend of weaker amplitudes in Ribo-seq-only rhythmic genes was also evident at a transcriptome-wide scale, as the average F.C. amplitude of Ribo-seq-only rhythmic genes (0.24 ± 0.01, s.e.m., *n* = 385) was considerably lower than that of both rhythmic genes (0.36 ± 0.02, *n* = 63) (see Fig. 1E, violin plots, p < 0.0001, *t*-test).

In the scatter plots in Fig. 1E–F, genes were plotted along the *x*-axis according to their average daily Ribo-seq abundance to highlight abundantly expressed genes in the SCN. This analysis revealed that *Gad1* (Glutamate Decarboxylase 1, essential for GABA production) is one of the most highly expressed Ribo-seq-only rhythmic genes in the SCN (Fig. 1F). However, it is crucial to mention that the normalized F.C. amplitude of *Gad1* was ∼0.21 and the ratio of its peak expression at ZT4 to its nadir expression at ZT20 was ∼2.0 (Fig. 1D). These values were relatively lower than those of representative both rhythmic genes, such as *Rgs16* and *Per2*.

The normalized F.C. amplitude of *Rgs16* was ∼0.66 and the ratio of its peak expression at ZT4 to its nadir expression at ZT16 was ∼7.0. These data indicate that there exist Ribo-seq-only rhythmic genes in the SCN, such as *Gad1*; however, they exhibit only modest oscillation amplitudes compared to the robust high-amplitude rhythms of both rhythmic genes.

### Oscillation profiles of ribosome-bound transcripts in the liver and kidney

Are the amplitudes characteristic of Ribo-seq-only rhythmic genes also conserved in other tissues? To address this question, we similarly characterized liver and kidney Ribo-seq-only rhythmic genes using publicly available datasets (GSE67305 for liver^17^ and GSE81283 for kidney^19^, Gene Expression Omnibus) (Fig. 2). We identified 544 and 390 Ribo-seq-only rhythmic genes in the liver and kidney, respectively (Supplementary Fig. 3, liver; Supplementary Fig. 4, kidney) and compared their oscillation amplitudes with the corresponding tissue’s both rhythmic genes. As exemplified by core clock genes such as *Per2*, *Bmal1*, and *Cry1*, transcripts that oscillated in both Ribo-seq and RNA-seq displayed robust oscillation amplitudes in the liver and kidney (see Supplementary Figs. 3 and 4)^17,19^. By contrast, as observed in the SCN, Ribo-seq-only rhythmic genes that were identified in the liver and kidney displayed less robust amplitudes, compared to both rhythmic genes (Fig. 2A for liver and Fig. 2B for kidney, violin plots, p < 0.0001, *t*-test).

**Figure 2.**
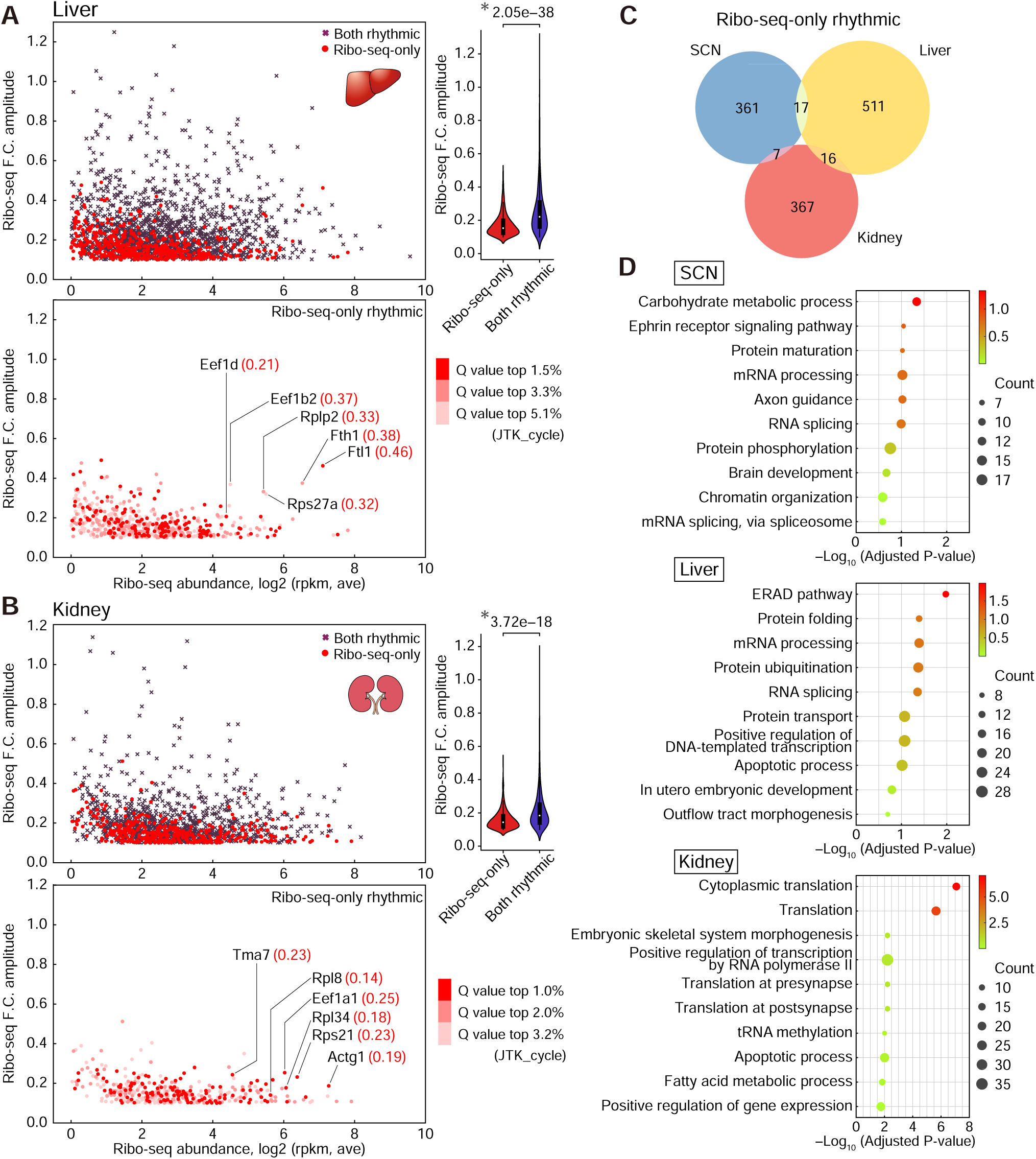
Tissue-specific and common features of Ribo-seq-only rhythmic genes identified by comparison between the SCN, liver, and kidney. (**A**–**B**) Scatter plots and violin plots for the liver (**A**) and kidney (**B**), plotted as described in Fig. 1E–F. Violin plots indicate significantly shallower amplitudes of Ribo-seq-only rhythmic genes than both rhythmic genes in the liver and kidney. The genes highlighted with F.C. amplitude values (red) are representative Ribo-seq-only rhythmic genes whose expression profiles are available in Supplementary Figs. 3 and 4. (**C**) Venn diagrams showing the overlap of SCN, liver, and kidney Ribo-seq-only rhythmic genes. (**D**) The top 10 enriched GO biological process terms of Ribo-seq-only rhythmic genes in the SCN, liver, and kidney. Statistics in (A) and (B), two-tailed *t*-test.

Not surprisingly, Ribo-seq-only rhythmic genes were highly tissue-specific, with no transcripts shared among the SCN, liver, and kidney (Venn diagram, Fig. 2C), suggesting that translational control may contribute to diversifying tissue-specific group of output genes^17,19^. Ribo-seq-only rhythmic genes in the SCN were enriched in those involved in brain functions (e.g. axon guidance and carbohydrate metabolism) (Fig. 2D, top), while in the liver and kidney they were enriched for posttranscriptional processes (e.g. ERAD pathway, protein folding and mRNA processing for the liver, and translation for the kidney), as has previously been reported^17–19^ (Fig. 2D, middle and bottom). These data suggest that the low-amplitude characteristic of Ribo-seq-only rhythmic genes is a common feature across tissue types, although the specific genes involved differ between tissues.

### Clock gene translation parallels mRNA rhythms with conserved efficiencies

As was observed in the liver and kidney^17,19^, circadian oscillations of core clock genes in the SCN predominantly occurred at the level of mRNA abundance (Fig. 3A) (see also liver and kidney clock genes in Supplementary Figs. 3 and 4). We observed that daily profiles of ribosome-bound transcripts (Ribo-seq counts, red lines in Fig. 3A) grossly correlate with those of their mRNA abundance (RNA-seq counts, blue lines in Fig. 3A) for all examined clock genes and clock-associated genes— *Per1*, *Per2*, *Per3*, *Cry1*, *Cry2*, *Bmal1*, *Clock*, *Npas2*, *Nr1d1*, *Nr1d2*, *Rora*, *Rorb*, *Dbp*, and *Nfil3*— in the SCN, which resemble those in the liver and kidney (Supplementary Figs. 5 and 6). However, one interesting exception was *Cry1*, which in the SCN —but not the liver or kidney — exhibited a slightly amplified Ribo-seq oscillation-amplitude compared with its RNA-seq rhythm, suggesting tissue-specific translational regulation of *Cry1*. On the other hand, no obvious translational regulation was detected for any clock genes; for example, no Ribo-seq-only rhythmic genes were found among clock genes (see also liver and kidney data in Supplementary Figs. 5 and 6).

**Figure 3.**
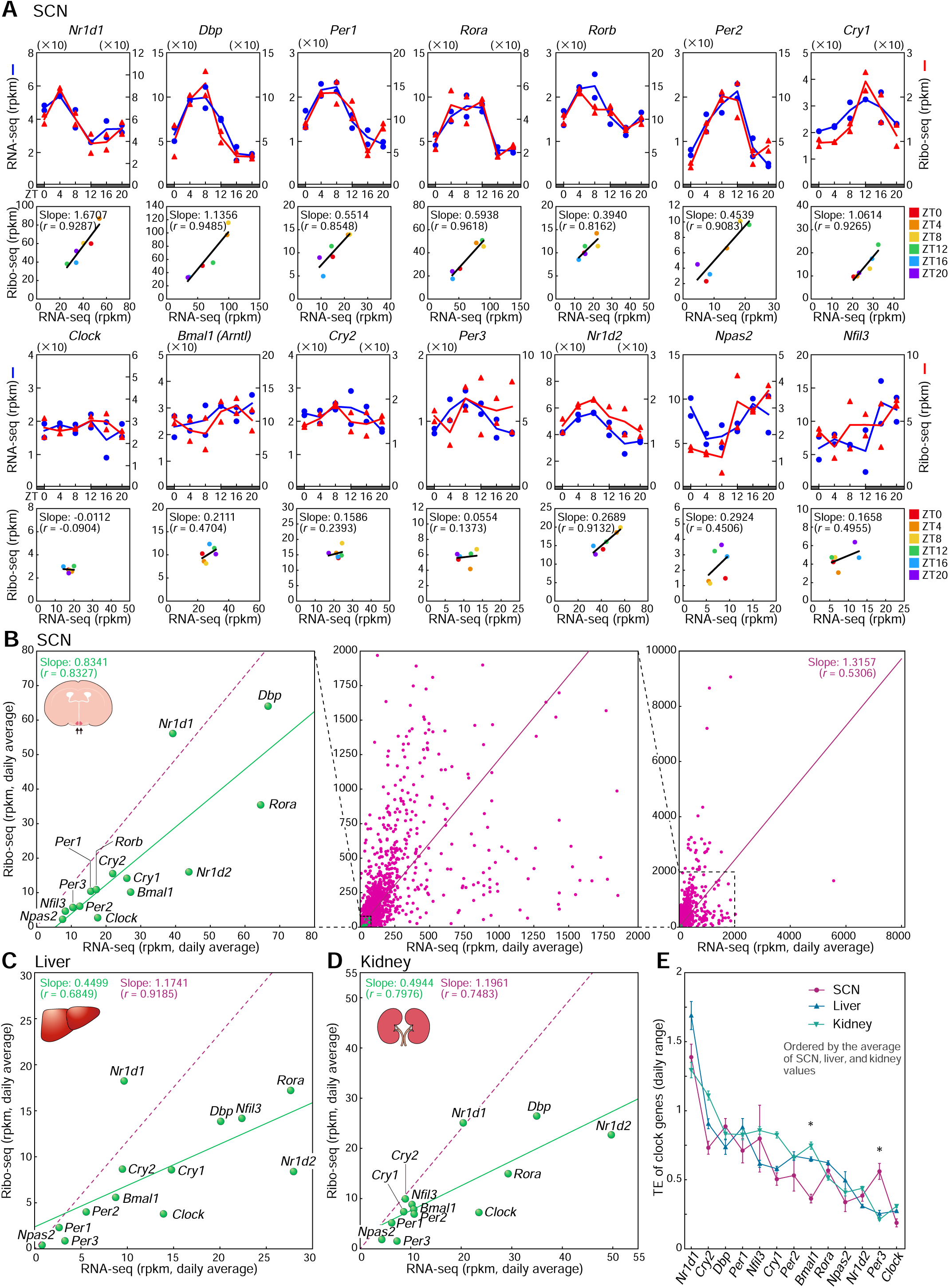
The proportion of bound ribosomes to RNA abundance in clock genes: near constant over time within each gene and heterogenous between genes. (**A**) Circadian RNA-seq and Ribo-seq profiles of 14 clock genes. Linear regression shows a positive correlation between the number of bound ribosomes and the abundance of RNA across ZT points tested within each gene. Colors denote different ZT points. (**B**) Transcriptome-wide scatter plots of SCN genes, plotted with RNA-seq values on the *x*-axis and Ribo-seq on the *y*-axis, illustrating the variety of the ratios of Ribo-seq to RNA-seq (i.e. translational efficiency) between clock genes (green) and among other expressed genes (red). (**C**–**D**) Scatter plots of clock genes in the liver (**C**) and kidney (**D**). The data are presented as described in (B). Transcriptome-wide plots and daily profiles of each clock gene in the liver and kidney are shown in Supplementary Figs. 5 and 6. (**E**) The TEs of clock genes in the SCN (red), liver (blue), and kidney (light green). Error bars represent SEM of TE values over the 24-hour timepoints. Asterisks indicate *Bmal1* and *Per3*; both displayed a greater than 2-fold significant difference in TE between the SCN and the liver and kidney. Statistics in (E), one-way ANOVA with Bonferroni post hoc test.

During the analysis of clock genes, we noticed considerable variation between clock genes in translation efficiency (TE), which is generally measured as the ratio of Ribo-seq footprint counts to RNA-seq read counts^16,21^. Because each clock gene did not exhibit notable TE variation across 24-h cycle^17–19^, we evaluated overall TE values for each clock gene using the 24-h average of the Ribo-seq and the corresponding RNA-seq data. We also plotted these values on a transcriptome-wide scale, with the daily average of the RNA-seq values on the *x*-axis and the corresponding Ribo-seq values on the *y*-axis (Fig. 3B). In this graph, the position of each plot (coordinates) reflects the TE value. For example, genes in the lower-right corner, such as *Nr1d2* and *Rora*, had relatively high mRNA abundance and modest ribosome binding. In contrast, genes located in the upper-left, such as *Nr1d1*, had relatively low mRNA abundance and higher ribosome binding, indicating a difference in TE between the clock genes. We also noted from the graph that clock genes were all expressed at relatively low levels compared to other abundantly expressed gene transcripts in the SCN, as has previously been reported^22^ (see Fig. 3B; compare clock genes marked in green versus non-clock-genes marked in red; clock genes were clustered in the lower-left corner of the transcriptome-wide scatter plots). The non-clock genes also exhibit a wide range of TEs as evident from their dispersed distribution across the plot.

Importantly, the same *x*–*y* plots of clock-gene transcripts in the liver and kidney resulted in graphs with a distribution pattern of clock genes roughly similar to what we observed in the SCN (Fig. 3C, liver; Fig. 3D, kidney). For example, *Nr1d1* was found to locate in the upper-left corner, while *Clock* and *Per2* in the bottom-left and *Nr1d2* in the middle-right, for the liver and kidney as well as the SCN. This suggests to us that although there is considerable variation in TE between clock genes, TEs of individual clock genes may be conserved across different tissues. To explore this possibility, we calculated the daily TE for all clock genes in the SCN, liver, and kidney, and permuted them based on the order of the TE means. The results showed that TE values varied from 0.25 to 1.5 depending on the type of clock genes, and that TE values for individual genes were overall comparable across tissues (see Fig. 3E; SCN, red; liver, blue; kidney, green). Statistical comparison also supported this conclusion, as a greater than twofold difference with significance (one-way ANOVA followed by Bonferroni’s test, p < 0.05) was only observed for *Bmal1* and *Per3* in comparisons of the SCN versus the liver and kidney (Fig. 3E, asterisks). Taken together, these data demonstrate that even in the SCN, mRNAs of all clock gene-components, either rhythmic or non-rhythmic, are translated largely concomitant with their accumulation, with TEs comparable to those observed in the liver and kidney.

### A screen for gene transcripts showing light-dependent ribosome binding

Our main interest in this study is to identify specific genes that are regulated or modulated at the level of mRNA translation. In this context, we next sought to obtain gene transcripts whose ribosome binding is modulated by light in the SCN. Light is the most potent environmental signal for entraining the central clock (and a signal uniquely processed by the SCN but not by peripheral tissue clocks). In our study, to screen for light-responsive translational regulation, mice were exposed to light for 4 h starting at ZT16, after which SCN tissues were collected and subjected to RNA-seq and Ribo-seq analyses. These light-at-night samples were generated as part of the same experimental series used for the daily time-course analysis shown in Fig. 1. Accordingly, the light-at-night data are presented alongside the daily expression profiles, with the ZT20 time point serving as the non-light control condition (Fig. 4A and Fig. 4B). In Fig. 4A, we present representative genes in which light exposure preferentially altered ribosome binding relative to mRNA abundance, whereas Fig. 4B highlights genes exhibiting parallel changes in ribosome binding and mRNA levels.

**Figure 4.**
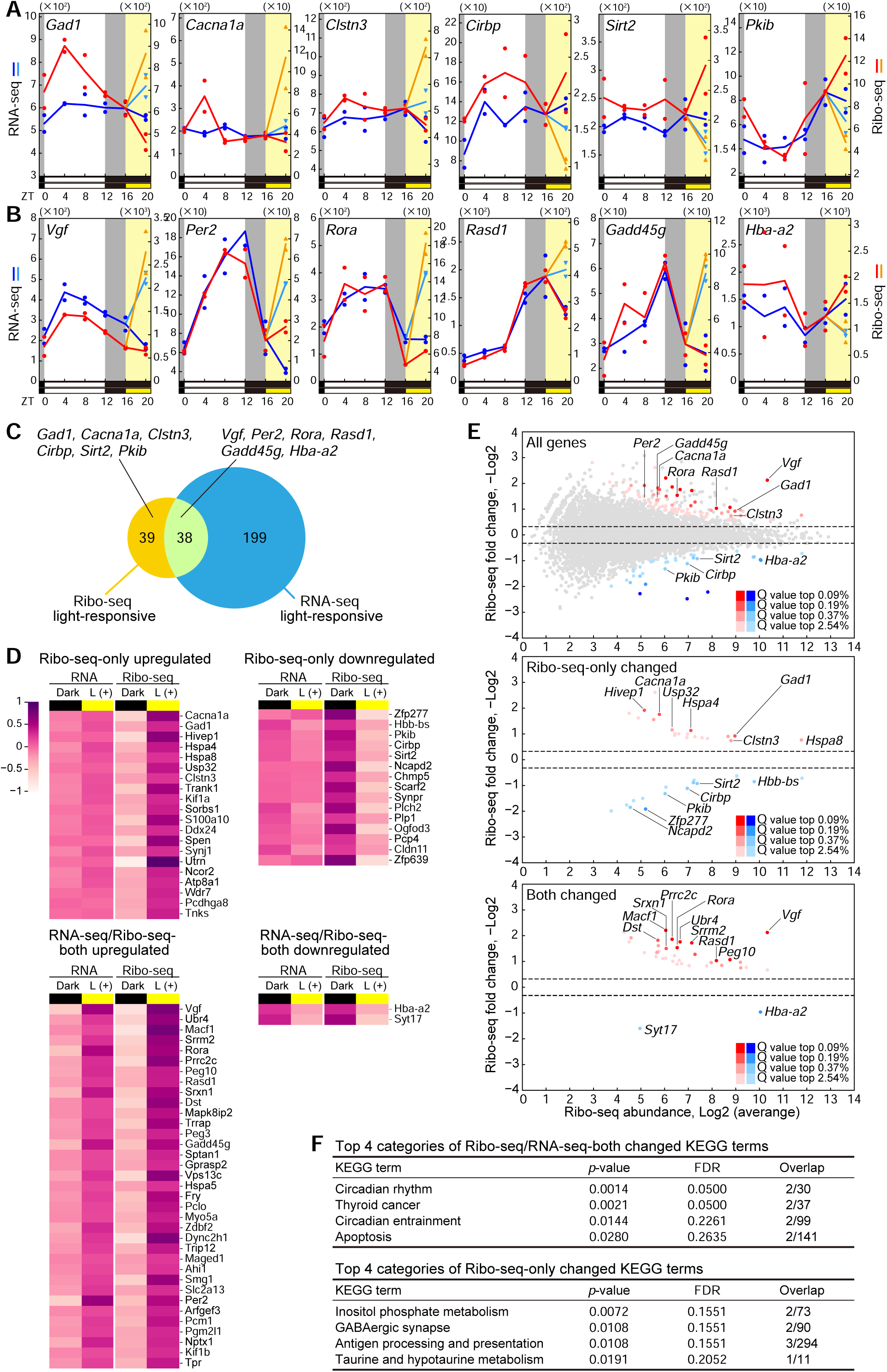
Light-regulated ribosome binding transcripts in the SCN. (**A**) Representative gene transcripts showing light-induced increase or decrease in ribosome binding. The SCN was sampled from mice after 4-h light illumination starting from ZT16. The data were overlaid on their daily Ribo-seq/RNA-seq profiles. The yellow shaded area represents time of light exposure. Orange and cyan lines show light-induced alterations in Ribo-seq and RNA-seq, respectively. The selected genes have a response to light greater in Ribo-seq than in RNA-seq. (**B**) Gene examples exhibiting light-induced changes in both Ribo-seq and RNA-seq. Data are shown as (A). (**C**) Venn diagram summarizing the number of genes light-regulated only in Ribo-seq, only in RNA-seq, and in both. Genes were considered as light-regulated if the change was more than 1.25-fold with a p value cutoff of 0.01 (DESeq2). (**D**) Heatmaps showing expression of genes upregulated or downregulated only in Ribo-seq and upregulated or downregulated in both Ribo-seq and RNA-seq. For color coding, the ratio to the geometric mean of the samples was calculated and log2 transformed. An RNA-seq fold change cutoff of 1.5 was applied for Ribo-seq-only genes. The gene order is based on the q-value for Ribo-seq change. (**E**) MA plots for all genes, Ribo-seq-only changed genes, and Ribo-seq/RNA-seq-both changed genes. The genes highlighted in the upper plots are those depicted in (A) and (B). The genes annotated in the middle and lower plots are q-value selected. (**F**) The top four KEGG terms of Ribo-seq/RNA-seq-both changed genes and Ribo-seq-only changed genes.

Data analysis on a transcriptome-wide scale (Fig. 4C) demonstrated that out of all detected transcripts in the SCN (11,172 transcripts, Fig. 1A), 77 responded at the ribosome binding level and 237 responded at the mRNA level, with an overlap of 38, when transcripts were considered light-responsive if they showed an increase or decrease of more than 1.25-fold with a p-value cutoff of 0.01 in Ribo-seq and RNA-seq (calculated using DESeq2). Genes of the following 4 categories, i) Ribo-seq-only upregulated, ii) Ribo-seq-only downregulated, iii) Ribo-seq/RNA-seq-both upregulated, and iv) Ribo-seq/RNA-seq-both downregulated, were individually illustrated in heatmaps (Fig. 4D). Genes were also plotted in MA-plot format (Fig. 4E) to emphasize their expression abundance, shown on the *x*-axis. These plots led us to notice that *Gad1* is one of the most highly expressed Ribo-seq-only light-inducible genes in the SCN (Fig. 4E, middle). As for the genes upregulated in both Ribo-seq and RNA-seq, *Vgf* (VGF nerve growth factor inducible) was among the most abundantly expressed genes with a relatively high fold change (Fig. 4E, bottom). Functional enrichment analysis revealed that genes showing light responsiveness in both Ribo-seq and RNA-seq were significantly enriched in KEGG pathways including “circadian rhythm” (FDR ≤ 0.05, Fig. 4F). On the other hand, for genes that changed only in Ribo-seq, no statistically significant enrichment was detected, while ‘‘GABA-ergic synapse’’ was highlighted as the second-highest KEGG term (FDR > 0.15).

Among the light-responsive transcripts identified, several genes attracted our attention based on their characteristic expression profiles shown in Fig. 4A and Fig 4B. For example, *Gad1* exhibited a more pronounced increase in ribosome binding than in mRNA abundance following light exposure; ribosome-bound *Gad1* transcripts increased by approximately twofold, reaching levels comparable to the daily peak observed at ZT4, whereas its mRNA levels showed only a modest increase, consistent with previously reported SCN microarray and RNA-seq data^23,24^. Likewise, light-dependent translational upregulation was observed for *Cacna1a* (Calcium Voltage-Gated Channel Subunit Alpha1 A) and *Clstn3* (Calsyntenin 3, synaptic regulator). On the other hand, downregulated translation was observed for *Cirbp* (Cold-inducible RNA Binding Protein), *Sirt2* (Sirtuin 2), and *Pkib* (Protein Kinase Inhibitor Beta). For instance, *Cirbp* showed a marked reduction in ribosome binding following light exposure (54%), whereas its mRNA levels were only modestly decreased (17%) compared with the non-light control (Fig. 4A). We also identified transcripts that were coordinately light-inducible in both Ribo-seq and RNA-seq, including the core clock genes *Per2* and *Rora*, as well as *Vgf*, all of which have been reported to increase in response to light in previous studies and SCN RNA-seq data^23–26^ (Fig. 4B). Thus, our data support previous findings and further demonstrated an increase in ribosome-bound transcripts for these genes in response to light.

### Gad1 protein product profiles in the SCN

Given that Ribo-seq reflects protein biosynthesis but not steady-state protein abundance, it is crucial to test whether the Ribo-seq changes we observed are accompanied by corresponding protein product accumulation in the SCN. We therefore examined protein abundance in the SCN by western blot analysis. Using pooled mouse SCN lysates (*n* = 6 punches per sample), the *Gad1*-encoded protein product Gad67 was readily detected, whereas western blot detection of other candidate proteins, including Cacna1a, Clstn3, Cirbp, Sirt2, and Pkib, was not successful under our experimental conditions, likely reflecting their low abundance in the SCN. Given that endogenous protein levels are relatively low, this poses challenges for immunodetection, both in terms of sensitivity and the potential for confounding nonspecific immunoreactive signals. In our protein blots, Gad67 was detected in SCN lysates as a single, discrete band migrating at ∼67 kDa, precisely matching the electrophoretic mobility of a recombinant Gad67 protein used as a reference control (Fig. 5A); no additional bands were observed, indicating high specificity of the signal detected in the SCN lysate. The same antibody also labeled brain regions consistent with reported Gad67 expression, with prominent enrichment in the SCN (Fig. 5B). This robust detection of Gad67 accords with the established neurochemical organization of the SCN, in which GABA serves as the principal neurotransmitter and almost all SCN neurons synthesize GABA (GABAergic)^7^.

**Figure 5.**
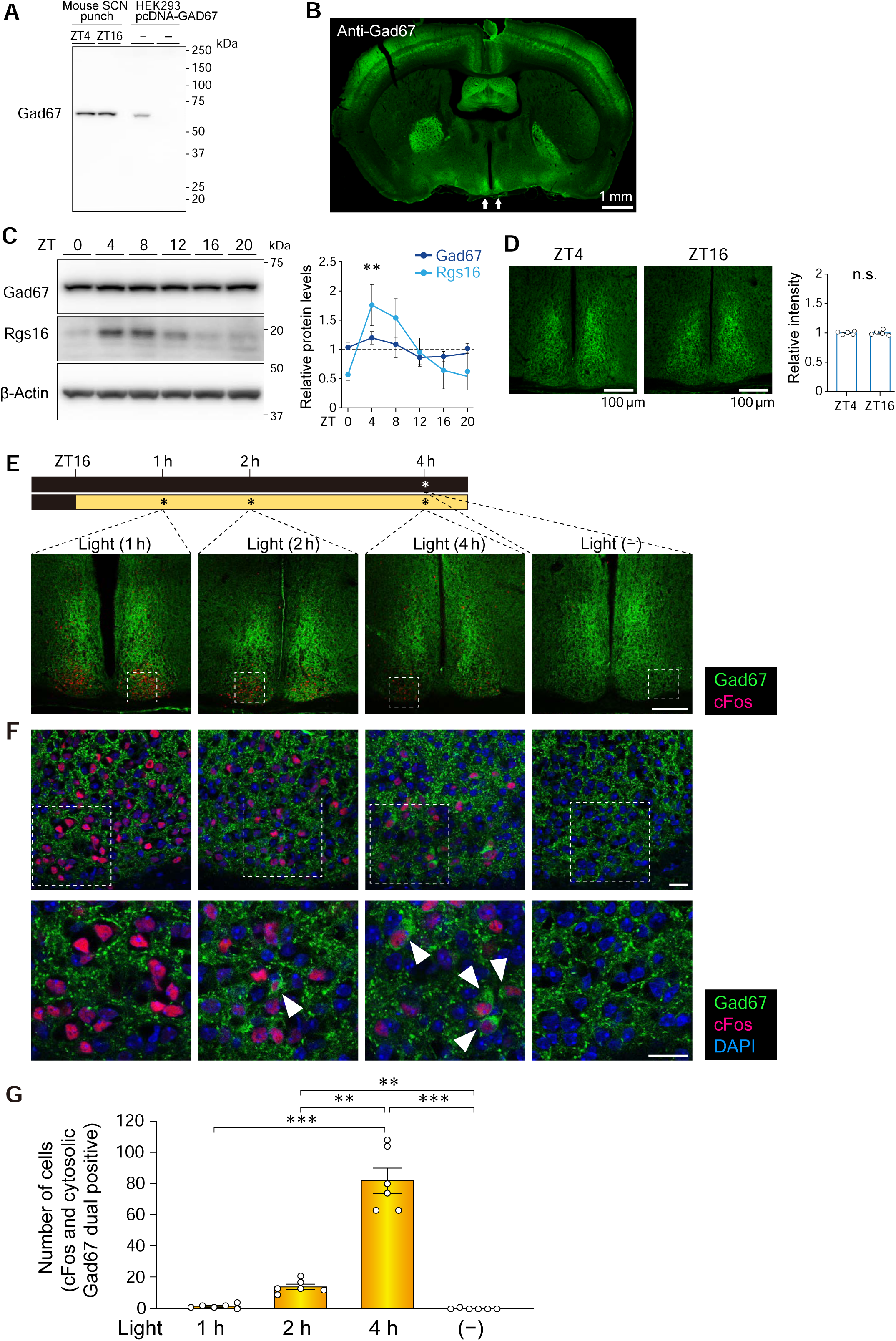
Increased cytosolic accumulation of Gad67 in light activated neurons in the SCN. (**A**) Uncropped western blots of Gad67. In SCN lysates, Gad67 was detected as a 67 kDa protein band, similar to the recombinant mouse Gad67 protein expressed in HEK293 cells. (**B**) A representative coronal brain section immunolabeled for Gad67. Arrows, the positions of the SCN. (**C**) Immunoblots of Gad67 at ZT0, 4, 8, 12, 16, 20 in the SCN. Rgs16 was detected as a control for a rhythmically-expressed protein. Protein band intensities, normalized to β-Actin, are expressed relative to the daily average expression. *n* = 3 biologically independent SCN samples. Error bars indicate SD. **p < 0.01, for Rgs16, ZT4 vs. ZT0 and ZT4 vs. ZT20, one-way ANOVA with Bonferroni’s post hoc test. (**D**) Anti-Gad67 SCN immunofluorescence staining, comparable between ZT4 and ZT16. (**E**) Immunofluorescence-based examination of Gad67 expression in mice exposed to light from ZT16 and sacrificed 1, 2, 4 h after light exposure. No-light-exposed animals at ZT20 served as control groups. (**F**) Representative coronal SCN sections immunolabeled for Gad67 (green) and cFos (red). Nuclei were stained with DAPI (blue). The dotted boxes indicate the regions enlarged in the bottom panels. Arrows point to cells displaying cytosolic Gad67 and nuclear cFos expression. Scale bars in (E) (F), 100 μm. (**G**) Quantification of Gad67/cFos double-positive cells in (F). The numbers of cells displaying cytosolic Gad67 and nuclear cFos expression were counted from the rostral to caudal extremities of the SCN. *n* = 6 SCN per group. Data are the mean ± SEM. Statistics, one-way ANOVA with Bonferroni post hoc test; **p < 0.01, ***p < 0.001, n.s., not significant.

Under these conditions, time-series western blot analysis revealed that Gad67 protein levels did not exhibit significant day–night variation in the SCN (see Fig. 5C). Gad67 protein band intensities (normalized with β-actin) had only shallow variations with no significance (Fig. 5C, right graph; *n* = 3 biologically independent SCN samples per time point). In marked contrast, the protein levels of the regulator of G-protein signaling 16 (Rgs16), which has been reported to exhibit robust daily rhythmic expression^27^, showed a pronounced day–night alteration at the protein level, with an approximately 3.5-fold increase peaking at ZT4 (p < 0.01, vs ZT0 and vs ZT20, see Fig. 5C). Notably, this peak phase coincided with the timing of maximal *Rgs16* ribosome binding observed in our Ribo-seq analysis (Fig. 1), validating the sensitivity of our immunoblot assay to detect daily protein oscillations. Immunohistochemical analysis likewise revealed no obvious difference in Gad67 immunofluorescence intensity between ZT4 and ZT16, in the SCN (Fig. 5D). Taken together, our results suggest that daily modulation of *Gad1* translation does not translate into overt day–night changes in steady-state Gad67 protein levels in the SCN.

For the purpose of scrutinizing Gad67 protein expression in response to light, we employed immunohistochemical examination of the SCN. More specifically, in this experiment, we scrutinized the potential light-dependent elevation of Gad67 protein immunofluorescence among cFos-positive neurons in the SCN (Fig. 5E–G); cFos was used, here, as a marker for staining light-activated SCN neurons according to previous reports^28–30^. Moreover, a time-course analysis was performed by examining Gad67 immunofluorescence at 1 h, 2 h, and 4 h after nocturnal light exposure (see a time scheme in Fig. 5E).

As expected, cFos immunofluorescence (*red*) was robustly induced within 1 h after nocturnal light exposure, consistent with the rapid light-dependent increase in *cFos* mRNA expression in the SCN (see Fig. 5E,F for protein, Supplementary Fig. 7A for mRNA qRT-PCR). This rapid transcriptional induction is in agreement with numerous previous reports^28,31^ and was further supported by reproduced SCN RNA-seq datasets (Supplementary Fig. 7B). In contrast, *Gad1* mRNA levels did not show a significant increase at either 1 h or 2 h following light exposure (Supplementary Fig. 7A), consistent with previously reported time-course RNA-seq data for *Gad1* in the SCN (Supplementary Fig. 7B).

Under these conditions, elevated Gad67 protein accumulation was detected selectively within a subset of light-activated SCN neurons, identified by cFos expression, after 4 h of light stimulation (Fig. 5E,F; Light (4 h)). As reported previously, cFos-positive light-responsive neurons were located in the retino-recipient region of the ventral SCN^29^. Noticeably, in these cells, elevated Gad67 accumulation was observed in the cytoplasmic (or perinuclear) region of each cell (see white arrowheads in Fig. 5F), consistent with the notion of nascent translation of the proteins.

By contrast, under no-light (dark) steady-state conditions, Gad67 immunofluorescence was not ubiquitously cytosolic but rather distributed in speckle-like structures (Fig. 5F, see Light (–)). On the basis of this qualitative difference, we focused our subsequent analyses on cells exhibiting concomitant cytosolic Gad67 accumulation and nuclear cFos expression. Notably, the number of cells displaying cytosolic Gad67 and nuclear cFos expression significantly increased after 4 h of light stimulation, whereas only a small number of cells exhibited this pattern after 1 or 2h light exposure (Fig. 5E,F, fluorescence images). Specifically, the number of double-positive cells per SCN was 82.0 ± 18.0 after 4 h, compared with 1.7 ± 1.2 at 1 h, 14.2 ± 3.8 at 2 h, and 0.2 ± 0.4 under dark conditions — a difference that was both large in magnitude and statistically highly significant (Fig. 5G, e.g., p < 0.001, 4 h vs. 1 h and Light (–)).

Together, these observations indicate that light-induced Gad67 protein accumulation in the SCN occurs with delayed kinetics relative to cFos induction and is not preceded by a drastic increase in *Gad1* mRNA, suggesting that this response is mediated, at least in part, at the level of protein translation.

## DISCUSSION

In this study, we employed ribosome profiling to investigate time-of-day-dependent and light-dependent translational regulation in the SCN, a principal brain region that serves as the master clock in the body. Through time-series RNA-seq/Ribo-seq analysis, we identified 385 genes that exhibited statistically significant rhythmicity at the level of ribosome association without corresponding oscillations in mRNA abundance. Notably, after light stimulation, *Gad1* transcripts exhibited an approximately twofold increase in ribosome binding, whereas total mRNA levels of *Gad1* were only marginally increased. Immunofluorescence staining revealed a subset of SCN neurons that exhibit light-dependent elevation of Gad67 protein accumulation. Unlike *Gad1*, all canonical circadian clock genes that constitute the transcription/translation-based feedback loops (TTFLs) were mainly regulated at the mRNA abundance level, with no additional evident regulation at the mRNA-translation level. Moreover, we found that Ribo-seq-only rhythmic genes (thus, translationally regulated genes) exhibited modest oscillation amplitudes compared to the robust high-amplitude rhythms of transcriptionally regulated genes. Our data thus identified differential regulatory features of transcription- and translation-based gene expression programs that perhaps together contribute to diverse gene expression profiles in the SCN.

A central question motivating this study was whether gene expression in the SCN can be regulated at the level of mRNA translation and, if so, whether such regulation can be uncovered by an unbiased, genome-wide Ribo-seq approach. In the present study, we provide the first ribosome-profiling (Ribo-seq) dataset generated from the SCN, enabling a genome-wide assessment of translational regulation in this central circadian pacemaker. Importantly, by combining Ribo-seq-based screening with protein-level validation, we demonstrate that light-dependent changes in ribosome binding, identified for *Gad1*, give rise to appreciable changes in protein product abundance in the SCN. These data therefore expand our current understanding of the modes of gene expression regulation in the SCN.

What might be the physiological significance of light-dependent translational upregulation of *Gad1* in the SCN? In general, GABAergic signaling has been interpreted as being regulated at the level of synaptic release, with Gad67/Gad65-dependent GABA biosynthesis viewed as a constitutive process. Our data thus added an additional layer to this framework by showing that light can induce *Gad1* translation in a subset of activated SCN neurons, leading to detectable accumulation of Gad67 protein without a preceding large increase in *Gad1* mRNA. This delayed, translation-dependent increase suggests that *de novo* GABA synthetic capacity may itself be dynamically tuned over hours in response to photic input. Physiologically, it is known that the SCN employs GABA signaling to coordinate SCN neuronal responses to light. Evidence in the literature indicates that GABA participates in maintaining intercellular phase synchrony between SCN neurons^32,33^, coordinating daily wake-rest cycles^34^, and encoding environmental or seasonal changes in day length^35,36^. Application of GABA causes phase-shifts in SCN neurons^37^. GABA receptor antagonists block phase-shifts caused by light stimulation^38,39^, and GABA signaling is reported to facilitate resynchronization between the SCN shell and core regions after phase dissociation induced by phase-shifting light exposure^40^. Thus, the protein expression elevation of Gad67, the enzyme responsible for GABA production, that we observed, may be explained as a compensatory mechanism to restore the GABA pool consumed in activated neurons in the process of light entrainment.

An important and unresolved question in the field of translational control is how gene-specific upregulation of translation is achieved in response to defined physiological stimuli. While global changes in translation efficiency have been extensively studied, the mechanisms that enable selective enhancement of protein synthesis from a restricted set of mRNAs—such as *Gad1* in the SCN—remain poorly understood. Theoretically, microRNAs targeting *Gad1* mRNA may be involved in *Gad1*-specific control^41^. It is also conceivable that yet-unknown functional mRNA structures and elements within the *Gad1* transcript, such as pseudoknots, hairpins, RNA G-quadruplexes, upstream open reading frames (uORFs), and internal ribosomal entry sites may mediate gene specificity^42^. The upstream signaling events mediating light-dependent *Gad1* translation also remain to be clarified. It has been reported that light can activate the p42/44 mitogen-activated protein kinase pathway leading to activation (phosphorylation) of the eukaryotic translation initiation factor 4E (eIF4E), which consequently increases the abundance of Per2 proteins in the SCN^43^, while our Ribo-seq data suggest that upregulation of *Per2* expression occurs mainly at the transcriptional level, at least based on the Ribo-seq data acquired at 4 h after the start of light stimulation at ZT16 (see Fig. 4B).

Although purely speculative, alternative upstream events may include reduced GABA pools in neurons, perhaps reflected by empty vesicles etc., which may trigger signaling to induce Gad67 protein expression via a currently unknown mechanism. Understanding how light causes *Gad1* mRNA translation in the SCN remains a challenge for future study.

TTFLs are the core molecular architecture of the circadian clock, generating circadian rhythmicity of mRNA/protein expression of a set of core oscillatory genes such as *Per2*, *Cry1*, and *Bmal1*^44,45^. Intuitively, we had expected that translation may be a critical regulatory step for rhythmical clock gene expression in the loops. However, as already revealed by studies analyzing translational regulation of clock genes in peripheral tissues—the liver and kidney—our Ribo-seq data for the SCN translatome confirmed that clock genes are mainly regulated at the mRNA abundance level with no evident regulation at the translation level. Our data thus align with the notion that, even in the central clock, transcript abundance control is the major driving force of oscillation in TTFL-based rhythm generation mechanism. Notwithstanding this conjecture, *Cry1* exhibited a slightly amplified Ribo-seq oscillation amplitude relative to RNA-seq specifically in the SCN (Fig. 3A). In this regard, it is interesting to note that *Cry1* protein abundance has been demonstrated to serve as a state variable of the SCN clock, as demonstrated by a study of translational switching in the SCN^46^. In addition, regarding translation efficiency (TE), we had expected that “different TEs” may contribute to cross-tissue differences in clock gene oscillations. However, we found that TEs of individual clock genes did not considerably vary between tissues, although there exist a few minor tissue-dependent difference in TE for *Bmal1* and *Per3* between the SCN and the liver or kidney (Fig. 3E). Our data thus suggest that individual TEs are conserved (fixed, gene by gene) across tissues, at least under steady-state circadian oscillation conditions. The functional meaning of distinct TEs for *Bmal1* and *Per3* is unknown and will be a subject of our future study. Overall, our data reinforce the importance of mRNA level regulation of clock genes.

With respect to oscillation amplitude, we found that Ribo-seq-only rhythmic genes share the characteristic of modest amplitudes, compared to the robust rhythms of transcriptionally-driven canonical clock genes. Crucially, this low-amplitude feature was also conserved in the liver and kidney Ribo-seq-only rhythmic genes (Fig. 2). Previous Ribo-seq studies analyzing liver and kidney circadian translatomes^17–19^ did not explicitly address this issue, but this low-amplitude characteristic is evident in their data. For example, representative Ribo-seq-only rhythmic genes in the liver that are reported by Janich *et al*.^17^ show modest normalized amplitudes. The amplitude values (JTK_cycle-determined amplitude divided by mean Ribo-seq expression) of genes reported for the liver, *Rps27a* (ribosomal protein S27a), *Fth1* (ferritin heavy polypeptide 1), and *Ftl1* (ferritin light polypeptide 1), were calculated as 0.32, 0.38, and 0.46, respectively. Similarly, the genes reported for the kidney by Castelo-Szekely *et al*.^19^, *Tma7* (translational machinery-associated 7), *Actg1* (actin gamma cytoplasmic 1), and *Hoxd3* (homeobox D3), exhibited amplitudes of 0.23, 0.19, and 0.31, respectively. These values were significantly smaller than those of the high-amplitude clock genes, e.g. *Per2* (0.94 for liver, 0.74 for kidney), *Cry1* (0.74 for liver, 0.79 for kidney), and *Bmal1* (0.97 for liver, 0.98 for kidney) (refs^17,19^; see also reanalyzed data in Supplementary Figs 3 and 4). One interpretation of this low-amplitude feature is that rhythms generated at the level of translation may have lesser biological significance. Yet, it is also equally plausible that low-amplitude rhythmicity at the level of translation could play a role in generating adaptive, subtle or fine-tuned adjustments of gene expression that cannot be achieved by transcriptional regulation alone. It is also worth noting that translational regulation can achieve (nucleus-independent) local translation inside the cell; this may be an additional merit of translational regulation, perhaps underlying *Gad1*-Gad67 regulation in the SCN.

Because Ribo-seq does not account for protein degradation or half-lives, our Ribo-seq data should not be interpreted as reflecting the abundance (steady-state level) of protein products in the SCN. Although our Ribo-seq data imply rhythmic synthesis of Gad67 in the SCN, protein blot data revealed that steady-state Gad67 protein levels in the SCN were nearly constant across the 24-h cycle. This constancy may result from a dynamic circadian equilibrium, where protein biosynthesis is balanced by degradation, or from a masking effect caused by a preexisting pool of accumulated proteins in the SCN. The same consideration is equally relevant when assessing other genes. For example, *Cirbp* and *Sirt2* caught our attention as genes whose translation levels were downregulated in response to 4 h light exposure (Fig. 4A). *Cirbp* is a potential mediator of circadian entrainment^47^, and *Sirt2* has been reported to mediate circadian rhythms of locomotor activity and energy expenditure^48^. However, our attempt at immunodetecting endogenous Cirbp and Sirt2 in the SCN failed because of the lack of appropriate antibodies for this purpose. The protein abundance of these genes, along with others identified in our Ribo-seq study, requires further investigation.

For summary, in the present study we have reported on the presence of translationally regulated genes involved in circadian clock, using the SCN as a model tissue. The SCN is the locus of the master circadian clock and constitutes the interface between external light and endogenous rhythms in the body. Previous studies of gene expression primarily focused on the transcriptome and proteome^13,23–25,49,50^, leaving translational regulation in the SCN unexplored. Therefore, the identification and demonstration of *Gad1* as the first example of an SCN light-regulated gene at the level of translation expand our current understanding of the modes of gene expression regulation in the SCN. A series of genome-wide transcriptome and translatome datasets generated for the SCN in this study may help to guide further investigation of translational control in the SCN.

## METHODS

### Animals

C57BL/6J male mice (6–8 wk old) were purchased from Japan SLC and entrained on a 12-h light/12-h dark cycle (lights on 8:00, lights off 20:00) with *ad libitum* access to food and water for at least 2 wk before experiments. In experiments examining light responsiveness, mice were exposed to light (∼200 lux) for 4 h from ZT16 (0:00). Animals were sacrificed by either cervical dislocation or transcardial perfusion-fixation at the indicated specific time-points. All tissue-sampling procedures in the dark period were performed under dim red light (>640 nm, <2 lux) conditions. All animal experiments in this study were performed in compliance with the Ethical Regulations of Kyoto University and conducted under protocols approved by the Animal Care and Experimentation Committee of Kyoto University.

### SCN punch

Microdissection of the SCN was performed as described^27^. In brief, mouse brains were quickly isolated from the skull and immediately frozen on dry ice. For each brain, a coronal brain section (300 μm thick) containing the SCN was prepared using a cryostat microtome (CM3050S, Leica) and mounted on a silicon rubber stage at −17 °C. Under a magnifying glass, the bilateral SCN was punched out from the frozen section using a 0.75 mm diameter punching needle (Leica Biosystems #39443001RM). The SCN punches from six animals per replicate were pooled together and lysed immediately in a specified buffer for ribosome profiling and western blotting (see below). The samples were stored at −80 °C until use.

### Ribo-seq and RNA-seq

For the time series analysis, we processed a total of 84 SCN punches (6 punches × 2 replicates × 7 time points, including one light illumination at night). The mouse SCN punches were lysed into ice-cold 20 mM Tris-HCl (pH 7.5) buffer containing 150 mM NaCl, 5 mM MgCl2, 1 mM dithiothreitol, 1% Triton X-100, and 100 μg ml^-1^ cycloheximide. After digesting genome DNA with DNase I (25 U ml^-1^, Thermo Fisher Scientific), lysates were centrifuged at 20,000 × g and the supernatant was processed for library generation as described^51,52^. For Ribo-seq, samples (0.5 μg RNA each) were digested with 20 U RNase (Epicentre) at 25 °C for 45 min, and the RNA fragments protected by ribosomes ranging 17–34 nt were gel-excised and ligated with linker oligonucleotides followed by rRNA depletion using Ribo-Zero Gold rRNA Removal Kit (Illumina). The resultant RNA fragments were reverse transcribed with Photoscript II (NEB) and circularized using CircLigaseII (Epicentre). The circularized cDNA templates were PCR amplified for 10 cycles using Phusion polymerase (NEB) and sequenced on an Illumina Hiseq X instrument. Footprints ranging from 20–34 nt were extracted and counted using the fp-count (https://github.com/ingolia-lab/RiboSeq). For RNA-seq (0.2 μg RNA each), samples were subjected to library construction using TruSeq Stranded Total RNA Gold (Illumina) and sequenced on HiSeq X as described^53^: Single-end reads (150 bp) were mapped onto the mouse genome (GRCm38/mm10) using STAR (version 2.7.0), sorted and indexed using SAMtools (version 1.10), and quantified using fp-count^54^. For filtering rRNA and tRNA, STAR was used with rRNA and tRNA annotations downloaded from the UCSC table browser.

### Bioinformatic for Ribo-seq and RNA-seq

Raw counts of the RNA-seq and Ribo-seq data were normalized using DESeq2. For genes with multiple isoforms, the most abundant isoform in RNA-seq was selected. For analysis of rhythmic genes, the nonparametric test JTK_cycle was used incorporating a period of 24 hours for the determination of circadian periodicity^20^. Output measures including amplitude and phase of rhythmic features were also determined. Genes were considered rhythmic in RNA-seq or Ribo-seq if their permutation-based, adjusted p-value (ADJ.P) was < 0.05 and further filtered using a JTK_cycle-determined amplitude cutoff of 0.1 (For the latter cutoff, values normalized using daily mean abundance were used; cf. Fig. 1D, schematic). In search for the genes that are rhythmic only in Ribo-seq or RNA-seq, we defined the genes as Ribo-seq-only rhythmic genes if Ribo-seq ADJ.P < 0.05 and RNA-seq ADJ.P > 0.2, and RNA-seq-only rhythmic genes if RNA-seq ADJ.P < 0.05 and Ribo-seq ADJ.P > 0.2. For comparison with RNA-seq and Ribo-seq public database from the mouse liver^17^ (GSE67305) and kidney^19^ (GSE81283), rhythmic transcripts were determined using a cutoff of BH.Q < 0.05 for both the liver and kidney and filtered using a JTK_cycle-determined amplitude cutoff of 0.1 as described in the SCN. Differential expression analysis in the SCN between ZT20 and 4 h-light groups was performed using DESeq2. Genes with p-values < 0.01 and fold change > 1.25 or < −1.25 were labeled as light-regulated genes. GO analysis was carried out using the DAVID bioinformatics resource^55^. KEGG pathway enrichment analysis was performed using GSEApy (https://github.com/ostrokach/gseapy)^56,57^. The complete lists of genes identified by RNA-seq and Ribo-seq are available in Supplementary Data: Supplementary Data 1 presents the SCN time-series RNA-seq and Ribo-seq analyses; Supplementary Data 2 and Supplementary Data 3 summarize the corresponding analyses for mouse liver and kidney, respectively; and Supplementary Data 4 compiles genes identified as light-responsive in the SCN.

### Western blotting

The SCN punches were pooled (*n* = 6 punches per sample) and directly lysed into Laemmli buffer containing 1x cOmplete Protease Inhibitor cocktail (Roche Diagnostics) as described^44^. Western blotting was performed according to our standard method using anti-Gad67 (mouse monoclonal, Millipore, MAB5406, 1:2,000 dilution), anti-Rgs16 (affinity-purified rabbit polyclonal^27^, 0.02 μg ml^-1^), anti-β-Actin (mouse monoclonal, A5441, Sigma-Aldrich, 1:1000), and anti-α-Tubulin (mouse monoclonal, Sigma Aldrich, T6199, 1:1,000) antibodies.

### Immunofluorescence

Free-floating immunofluorescence staining was performed using 30-μm-thick serial coronal brain sections using anti-Gad67 (mouse monoclonal, Millipore, MAB5406, 1:500) and anti-cFos (rabbit polyclonal, Abcam, ab190289, 1:500) for 24 h at 4 °C. Immunoreactivities were visualized with Alexa488-conjugated anti-mouse IgG (Thermo Fisher Scientific, 1:1,000) and Alexa594-conjugated anti-rabbit IgG (Thermo Fisher Scientific, 1:1,000). The sections were mounted in medium containing 4ʹ,6ʹ -diamidino-2-phenylindole (DAPI) for nuclear staining. For brightness and contrast, photomicrographs were processed identically with ImageJ. For quantitative analysis, serial coronal brain sections (30μm thick), covering from the rostralmost to the caudalmost of the SCN (20 sections per animal), were counted. To minimize technical variations in immunostaining, sections from different experimental conditions were gathered into one group and processed simultaneously as described^58^. An observer blind to the experimental condition counted the number of positive cells in the SCN.

## Statistics analysis

Western blot band and immunofluorescence intensities were quantified using Image J software. Statistical analysis was performed using GraphPad Prism 8 with the statistical tests shown in the figure legends.

## Data availability

Data that support the findings of this study are available as Source Data and Supplementary Data. RNA-seq datasets generated in this study are available at Gene Expression Omnibus (accession number: GSE315833).

## ACKNOWLEDGEMENTS

This work was supported in part by research grants from the Ministry of Education, Culture, Sports, Science and Technology of Japan (JP22H04987, JP24H02306, and JP24H02307), the Basis for Supporting Innovative Drug Discovery and Life Science Research program of the Japan Agency for Medical Research and Development (JP21am0101092), SRF, and Astellas Foundation for Research on Metabolic Disorders. X.S. is a recipient of the JSPS Research Fellowship for Young Scientists. Computation was supported by the HOKUSAI SailingShip supercomputer facility at RIKEN.

## Author Contributions

M.D. conceived the project; M.D. and X.S. designed the research; X.S. performed experiments in collaboration with S.-W. H., C.X., H.Z., Y.T., T.M., Y.S., S.I., and E.H; M.D. and X.S. wrote the paper with input from all authors.

## Declaration of interests

The authors declare no competing financial interests.

## Additional information

Supplementary information is available for this paper.

Correspondence and requests for materials should be addressed to M.D.

## Supplementary Information For

**Supplementary Figure 1 (related to Fig. 1).**
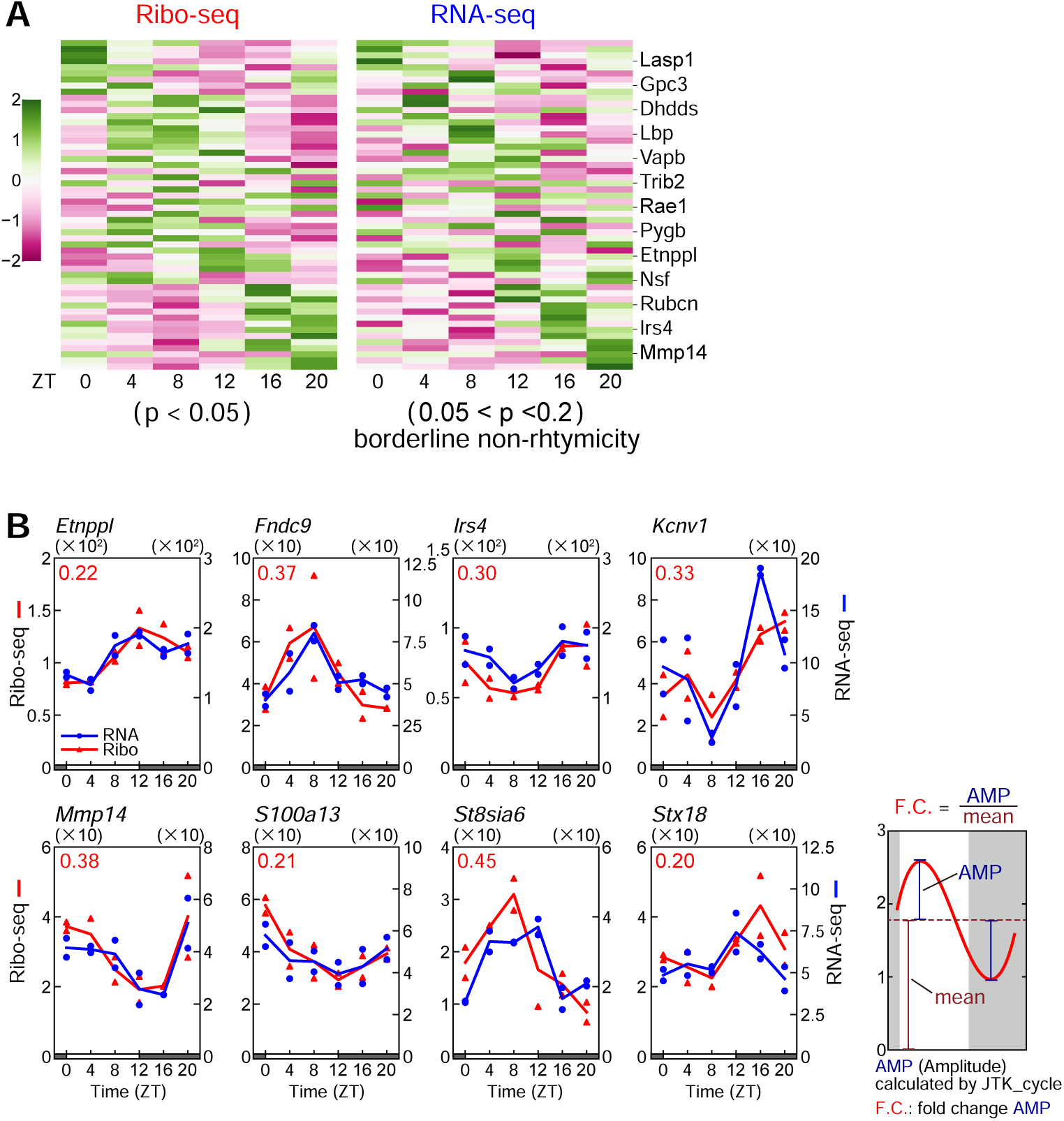
Expression profiles of genes displaying Ribo-seq rhythmicity (p < 0.05) and non-significant borderline RNA-seq rhythmicity (0.05 < p < 0.2) by JTK_cycle analysis. (A) Heatmaps of Ribo-seq and RNA-seq profiles. (B) Representative oscillation profiles of corresponding genes. Data were analyzed ad described in Fig. 1D.

**Supplementary Figure 2 (related to Fig. 1).**
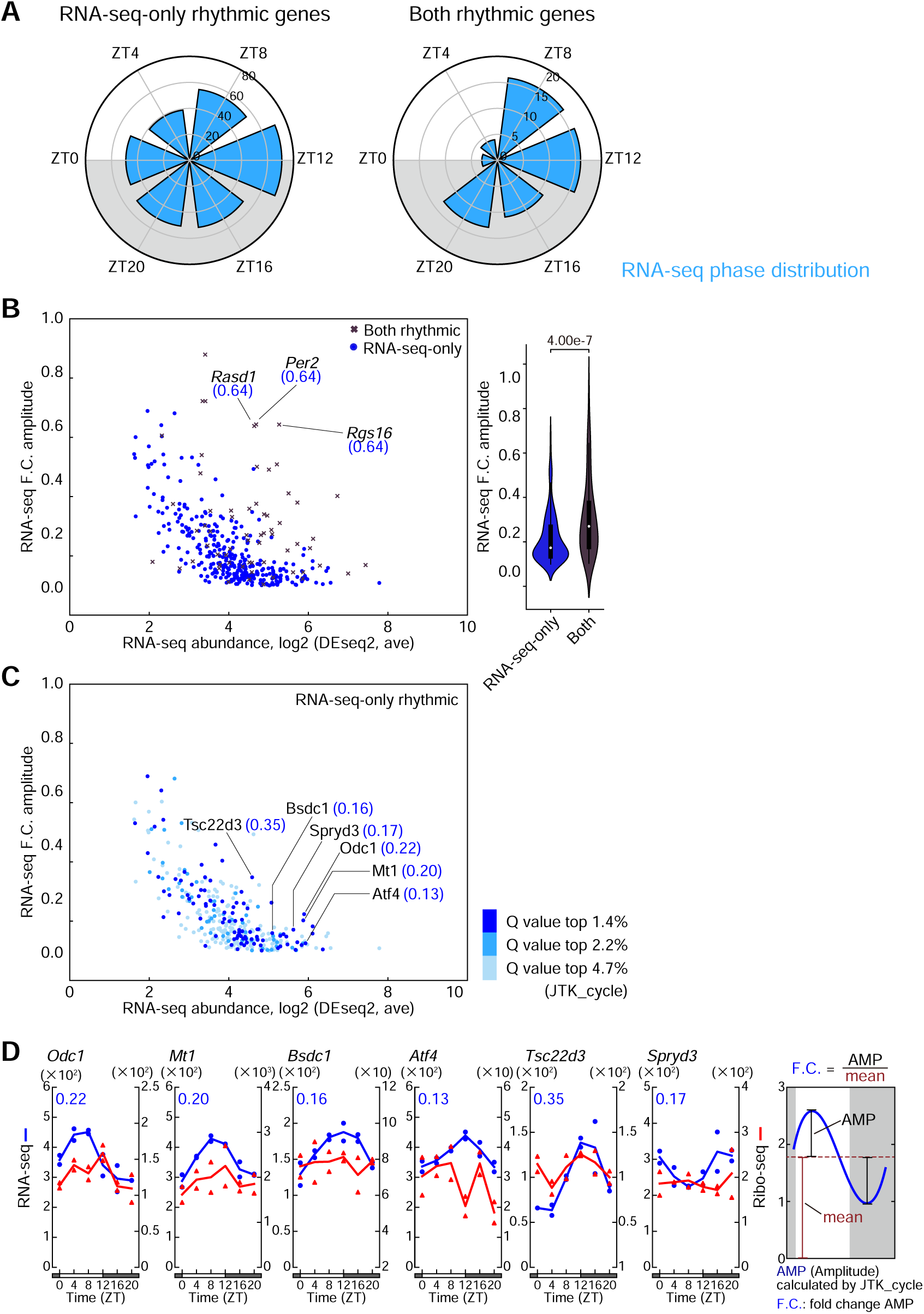
Transcripts displaying rhythmicity only in RNA-seq in the SCN. (A) Circular histogram showing the peak phase of RNA-seq specific genes and both rhythmic genes. (B) Scatter plots of both rhythmic, RNA-seq-only, and Ribo-seq-only genes, charting RNA-seq rhythm amplitude and abundance. Violin plots show the range of amplitude for each category. (C) Representative RNA-seq-only genes. The plots are the same as (B), with colors varying by q-value (JTK_cycle). (D) Circadian RNA-seq/Ribo-Seq profiles of genes highlighted in (C). The blue values on the graphs represent fold change amplitude (F.C.) of the RNA-seq rhythm, calculated as the ratio of the JTK_cycle determined amplitude (AMP) to mean RNA-seq abundance. Statistics in (B), two-tailed unpaired *t*-test.

**Supplementary Figure 3 (related to Fig. 2A).**
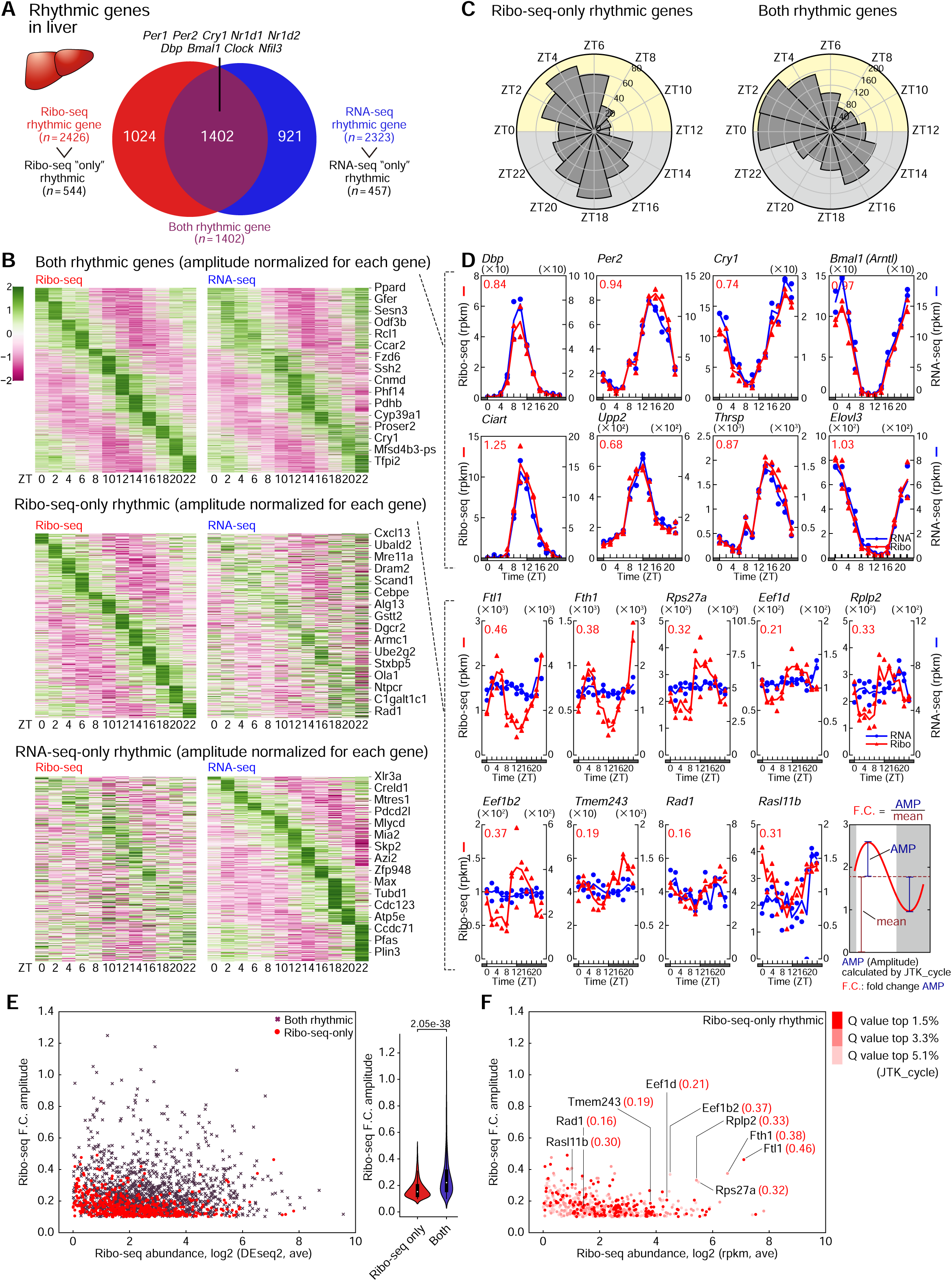
Identification of transcripts with circadian ribosome binding in the liver. (A) Venn diagram showing the number of Ribo-seq rhythmic genes (red) and RNA-seq rhythmic genes (blue). The overlap corresponds to both rhythmic genes. (B) Heatmaps of ribosome binding (left) and mRNA abundance (right) rhythms for both rhythmic genes (top), Ribo-seq-only rhythmic genes (middle), and RNA-seq-only rhythmic genes (bottom). Amplitudes were standardized within each gene using Z-scores. (C) Polar histogram plots of the peak phase for Ribo-seq-only rhythmic genes and both rhythmic genes. (D) Representative oscillation profiles of both rhythmic genes and Ribo-seq-only rhythmic genes. Ribo-seq-only genes were selected based on their lack of evident RNA-seq rhythmicity. The red values on the graphs represent F.C. amplitude of Ribo-seq rhythm, as described in Fig. 1D. (E) Scatter plots of both rhythmic and Ribo-seq-only rhythmic genes, plotted with daily average Ribo-seq abundance on the *x*-axis and Ribo-seq F.C. amplitude on the *y*-axis. Violin plots show the range of amplitude for each category. (F) Replots of Ribo-seq-only rhythmic genes. The genes highlighted with F.C. amplitude values are representative Ribo-seq-only rhythmic genes whose expression profiles are shown in (D). Statistics in (E), two-tailed unpaired *t*-test.

**Supplementary Figure 4 (related to Fig. 2B).**
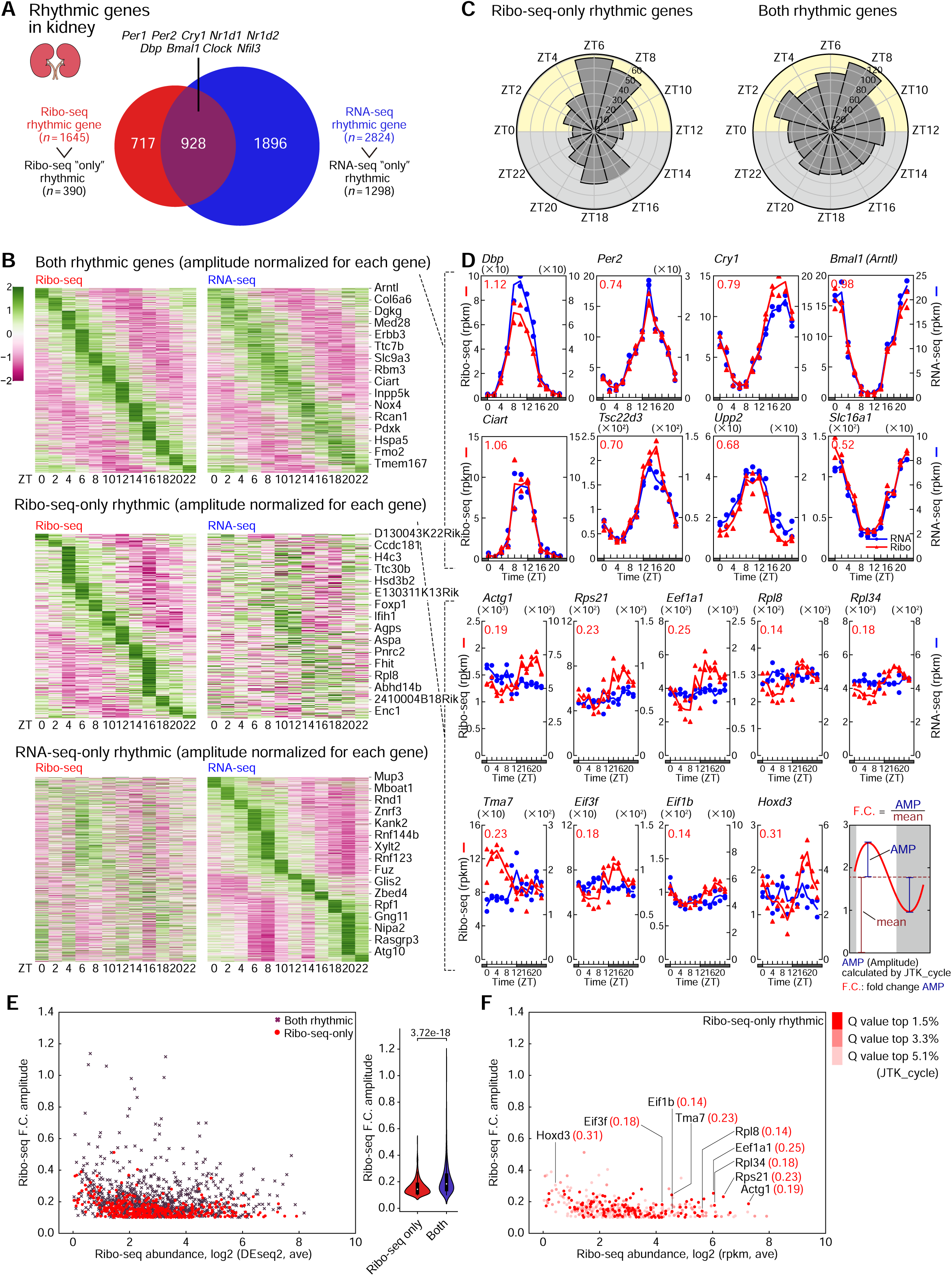
Identification of transcripts with circadian ribosome binding in the kidney. (A) Venn diagram showing the number of Ribo-seq rhythmic genes (red) and RNA-seq rhythmic genes (blue). The overlap corresponds to both rhythmic genes. (B) Heatmaps of ribosome binding (left) and mRNA abundance (right) rhythms for both rhythmic genes (top), Ribo-seq-only rhythmic genes (middle), and RNA-seq-only rhythmic genes (bottom). Amplitudes were standardized within each gene using Z-scores. (C) Polar histogram plots of the peak phase for Ribo-seq-only rhythmic genes and both rhythmic genes. (D) Representative oscillation profiles of both rhythmic genes and Ribo-seq-only rhythmic genes. Ribo-seq-only genes were selected based on their lack of evident RNA-seq rhythmicity. The red values on the graphs represent F.C. amplitude of Ribo-seq rhythm, as described in Fig. 1D. (E) Scatter plots of both rhythmic and Ribo-seq-only rhythmic genes, plotted with daily average Ribo-seq abundance on the *x*-axis and Ribo-seq F.C. amplitude on the *y*-axis. Violin plots show the range of amplitude for each category. (F) Replots of Ribo-seq-only rhythmic genes. The genes highlighted with F.C. amplitude values are representative Ribo-seq-only rhythmic genes whose expression profiles are shown in (D). Statistics in (E), two-tailed unpaired *t*-test.

**Supplementary Figure 5 (related to Fig. 3C).**
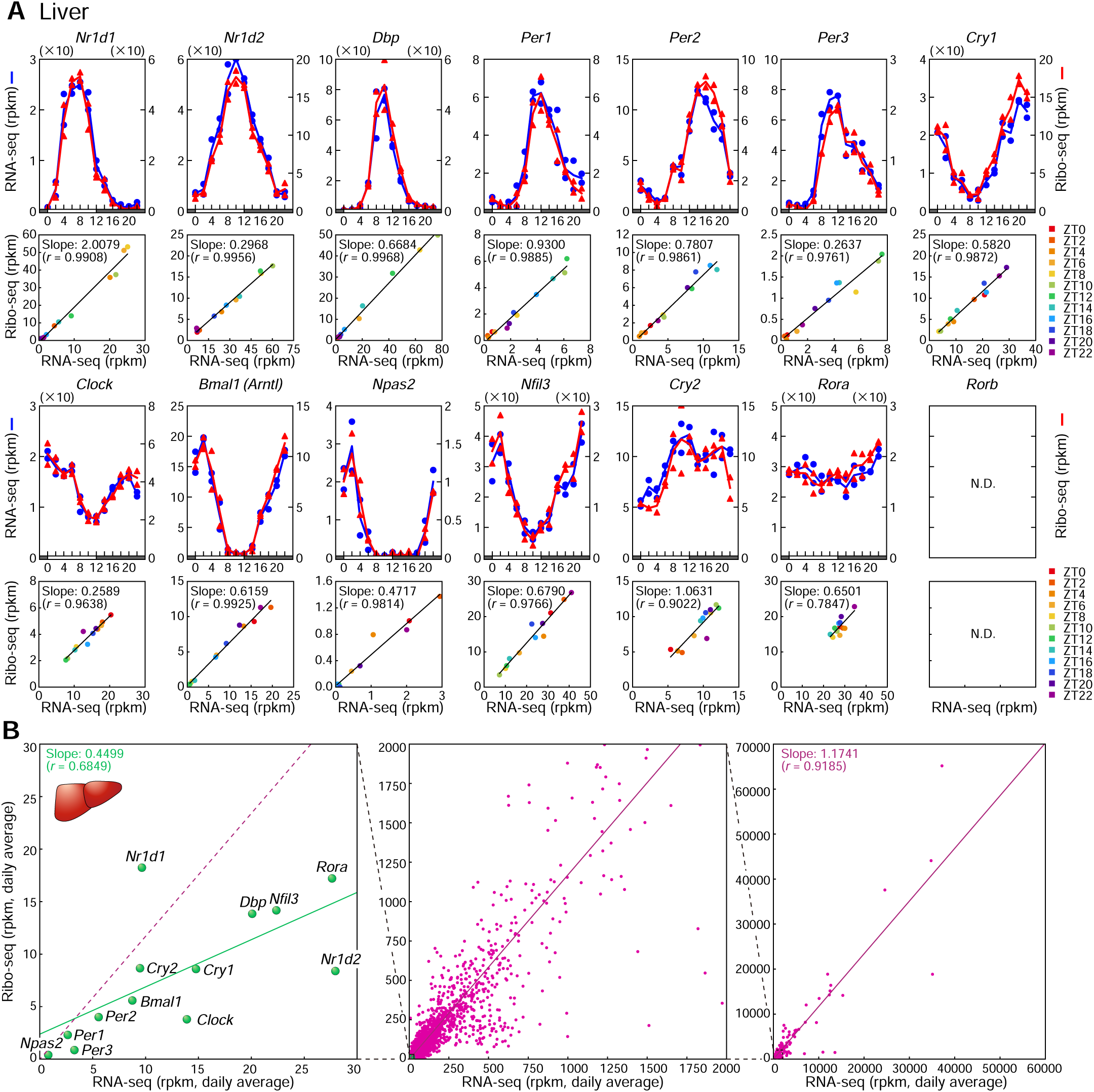
Highly correlated rhythms of ribosome binding and mRNA abundance of liver clock genes with translational efficiency variety between genes. (A) RNA-seq and Ribo-seq profiles of 14 clock genes in the liver. Colors denote different timepoints. Data were analyzed as described in Fig. 3A. N.D., not detected. (B) Transcriptome-wide scatter plots of liver genes, plotted with RNA-seq values on the *x*-axis and Ribo-seq on the *y*-axis, illustrating the variety of the ratios of Ribo-seq to RNA-seq (i.e. TEs) between clock genes (green) and among other expressed genes (red).

**Supplementary Figure 6 (related to Fig. 3D).**
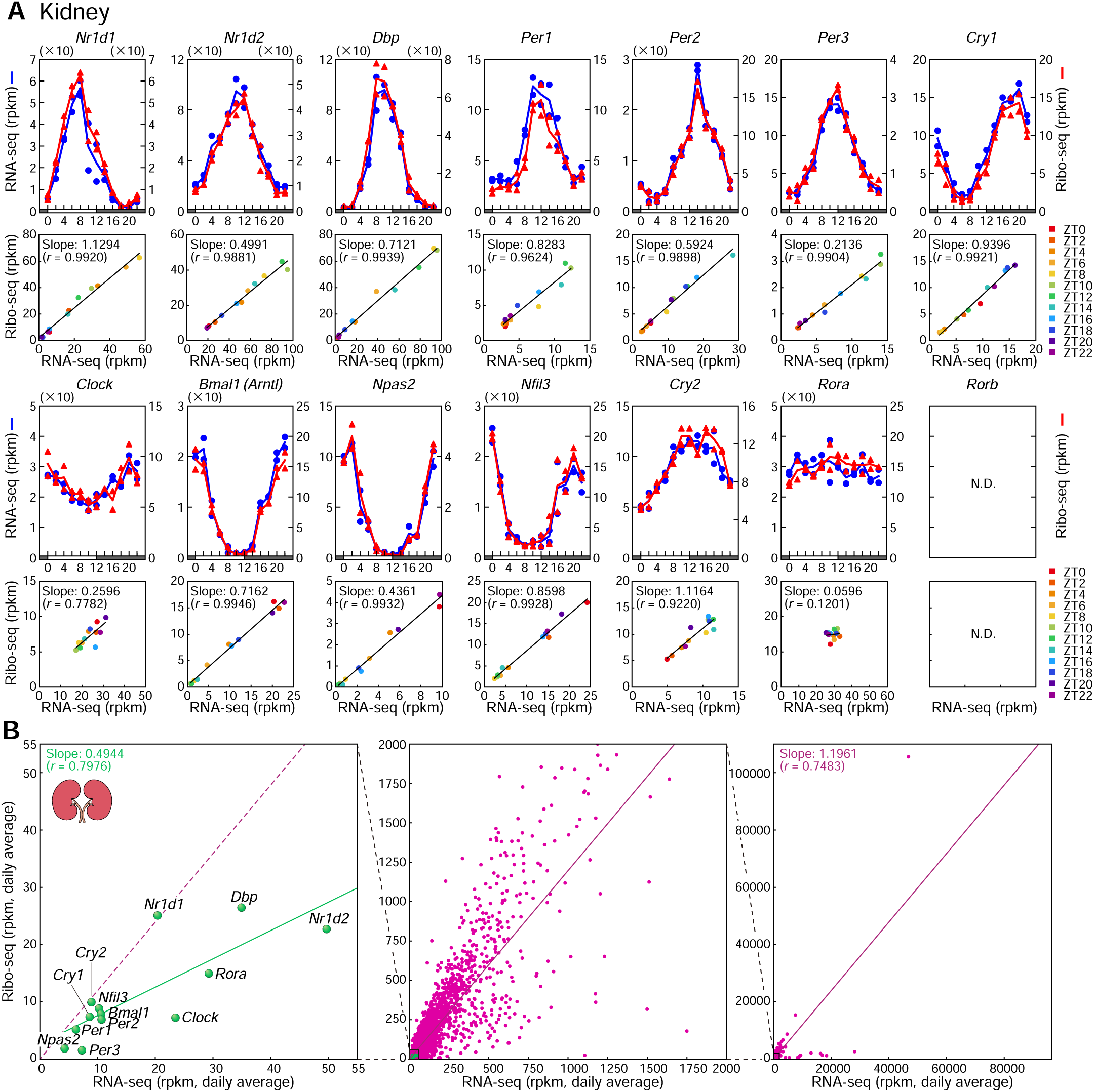
Highly correlated rhythms of ribosome binding and mRNA abundance of kidney clock genes with translational efficiency variety between genes. (A) RNA-seq and Ribo-seq profiles of 14 clock genes in the kidney. Colors denote different timepoints. Data were analyzed as described in Fig. 3A. N.D., not detected. (B) Transcriptome-wide scatter plots of kidney genes, plotted with RNA-seq values on the *x*-axis and Ribo-seq on the *y*-axis, illustrating the variety of the ratios of Ribo-seq to RNA-seq (i.e. TEs) between clock genes (green) and among other expressed genes (red).

**Supplementary Figure 7 (related to Fig. 5).**
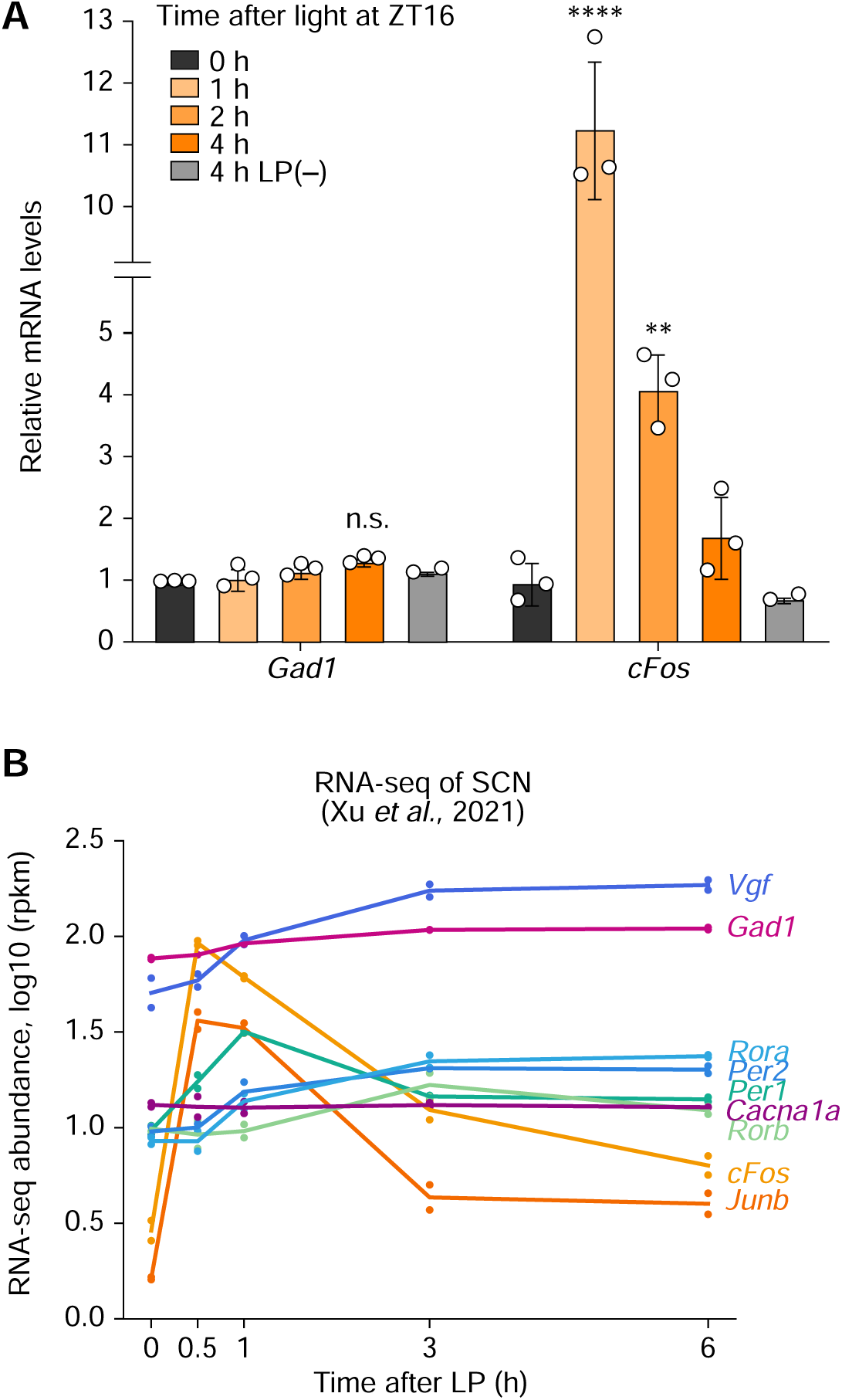
A modest but non-significant increase in mRNA abundance of *Gad1* after light exposure. (A) Expression profiles of *Gad1* and *cFos* mRNA in the SCN after light exposure at ZT16. Mice were sacrificed 0, 1, 2, 4 h after light stimulation. No-light-exposed animals at ZT20 (4 h LP (–)) served as control groups. The SCN was dissected out by laser microdissection, and mRNA levels of *Gad1* and *cFos* were determined by gene-specific qRT-PCR and normalized to *Rplp0*. *n* = 3 biologically independent SCN samples per data point. Data are plotted as mean ± SEM, relative to the basal values at ZT16. (B) Temporal profiles of representative light-responsive genes in the SCN, reproduced from RNA-seq data reported by Xu *et al*.^23^ Statistics in (A), **p < 0.01, for *cFos*, 2 h vs 0 h, ****p < 0.0001, for *cFos*, 1 h vs 0 h, n.s., not significant, for *Gad1*, 4 h vs 4 h LP (–), two-way ANOVA with Bonferroni multiple comparisons test.

## Notes

### Competing Interest Statement

The authors have declared no competing interest.

## REFERENCES

1. Mohawk, J.A., Green, C.B. & Takahashi, J.S. Central and peripheral circadian clocks in mammals. Annu Rev Neurosci 35, 445–462 (2012).

2. Herzog, E.D., Hermanstyne, T., Smyllie, N.J. & Hastings, M.H. Regulating the Suprachiasmatic Nucleus (SCN) Circadian Clockwork: Interplay between Cell-Autonomous and Circuit-Level Mechanisms. Cold Spring Harb Perspect Biol 9(2017).

3. Hastings, M.H., Maywood, E.S. & Brancaccio, M. Generation of circadian rhythms in the suprachiasmatic nucleus. Nat Rev Neurosci 19, 453–469 (2018).

4. Hsiao, S.W. & Doi, M. Circuits involving the hypothalamic suprachiasmatic nucleus for controlling diverse physiologies verified by the aid of optogenetics and chemogenetics. Int Rev Cell Mol Biol 393, 1–14 (2025).

5. Asada, H., et al. Cleft palate and decreased brain gamma-aminobutyric acid in mice lacking the 67-kDa isoform of glutamic acid decarboxylase. Proc Natl Acad Sci U S A 94, 6496–6499 (1997).

6. Kash, S.F., et al. Epilepsy in mice deficient in the 65-kDa isoform of glutamic acid decarboxylase. Proc Natl Acad Sci U S A 94, 14060–14065 (1997).

7. Moore, R.Y. & Speh, J.C. GABA is the principal neurotransmitter of the circadian system. Neurosci Lett 150, 112–116 (1993).

8. Koh, W., Kwak, H., Cheong, E. & Lee, C.J. GABA tone regulation and its cognitive functions in the brain. Nat Rev Neurosci 24, 523–539 (2023).

9. Patz, S., Wirth, M.J., Gorba, T., Klostermann, O. & Wahle, P. Neuronal activity and neurotrophic factors regulate GAD-65/67 mRNA and protein expression in organotypic cultures of rat visual cortex. Eur J Neurosci 18, 1–12 (2003).

10. Chattopadhyaya, B., et al. GAD67-mediated GABA synthesis and signaling regulate inhibitory synaptic innervation in the visual cortex. Neuron 54, 889–903 (2007).

11. Hanno-Iijima, Y., Tanaka, M. & Iijima, T. Activity-Dependent Bidirectional Regulation of GAD Expression in a Homeostatic Fashion Is Mediated by BDNF-Dependent and Independent Pathways. PLoS One 10, e0134296 (2015).

12. Pembroke, W.G., Babbs, A., Davies, K.E., Ponting, C.P. & Oliver, P.L. Temporal transcriptomics suggest that twin-peaking genes reset the clock. Elife 4(2015).

13. Panda, S., et al. Coordinated transcription of key pathways in the mouse by the circadian clock. Cell 109, 307–320 (2002).

14. Huhman, K.L., Hennessey, A.C. & Albers, H.E. Rhythms of glutamic acid decarboxylase mRNA in the suprachiasmatic nucleus. J Biol Rhythms 11, 311–316 (1996).

15. Ingolia, N.T. Ribosome profiling: new views of translation, from single codons to genome scale. Nat Rev Genet 15, 205–213 (2014).

16. Ingolia, N.T., Ghaemmaghami, S., Newman, J.R. & Weissman, J.S. Genome-wide analysis in vivo of translation with nucleotide resolution using ribosome profiling. Science 324, 218–223 (2009).

17. Janich, P., Arpat, A.B., Castelo-Szekely, V., Lopes, M. & Gatfield, D. Ribosome profiling reveals the rhythmic liver translatome and circadian clock regulation by upstream open reading frames. Genome Res 25, 1848–1859 (2015).

18. Atger, F., et al. Circadian and feeding rhythms differentially affect rhythmic mRNA transcription and translation in mouse liver. Proc Natl Acad Sci U S A 112, E6579–6588 (2015).

19. Castelo-Szekely, V., Arpat, A.B., Janich, P. & Gatfield, D. Translational contributions to tissue specificity in rhythmic and constitutive gene expression. Genome Biol 18, 116 (2017).

20. Hughes, M.E., Hogenesch, J.B. & Kornacker, K. JTK_CYCLE: an efficient nonparametric algorithm for detecting rhythmic components in genome-scale data sets. J Biol Rhythms 25, 372–380 (2010).

21. Ingolia, N.T., Lareau, L.F. & Weissman, J.S. Ribosome profiling of mouse embryonic stem cells reveals the complexity and dynamics of mammalian proteomes. Cell 147, 789–802 (2011).

22. Zhang, R., Lahens, N.F., Ballance, H.I., Hughes, M.E. & Hogenesch, J.B. A circadian gene expression atlas in mammals: implications for biology and medicine. Proc Natl Acad Sci U S A 111, 16219–16224 (2014).

23. Xu, P., et al. NPAS4 regulates the transcriptional response of the suprachiasmatic nucleus to light and circadian behavior. Neuron 109, 3268–3282.e3266 (2021).

24. Hatori, M., et al. Lhx1 maintains synchrony among circadian oscillator neurons of the SCN. Elife 3, e03357 (2014).

25. Wen, S., et al. Spatiotemporal single-cell analysis of gene expression in the mouse suprachiasmatic nucleus. Nat Neurosci 23, 456–467 (2020).

26. Okamura, H., et al. Photic induction of mPer1 and mPer2 in cry-deficient mice lacking a biological clock. Science 286, 2531–2534 (1999).

27. Doi, M., et al. Circadian regulation of intracellular G-protein signalling mediates intercellular synchrony and rhythmicity in the suprachiasmatic nucleus. Nat Commun 2, 327 (2011).

28. Rusak, B., Robertson, H.A., Wisden, W. & Hunt, S.P. Light pulses that shift rhythms induce gene expression in the suprachiasmatic nucleus. Science 248, 1237–1240 (1990).

29. Colwell, C.S. & Foster, R.G. Photic regulation of Fos-like immunoreactivity in the suprachiasmatic nucleus of the mouse. J Comp Neurol 324, 135–142 (1992).

30. Yamaguchi, Y., et al. Gpr19 is a circadian clock-controlled orphan GPCR with a role in modulating free-running period and light resetting capacity of the circadian clock. Sci Rep 11, 22406 (2021).

31. Kornhauser, J.M., Mayo, K.E. & Takahashi, J.S. Light, immediate-early genes, and circadian rhythms. Behav Genet 26, 221–240 (1996).

32. Bussi, I.L., et al. Expression of the vesicular GABA transporter within neuromedin S. Proc Natl Acad Sci U S A 120, e2314857120 (2023).

33. Ono, D., Honma, K.I., Yanagawa, Y., Yamanaka, A. & Honma, S. GABA in the suprachiasmatic nucleus refines circadian output rhythms in mice. Commun Biol 2, 232 (2019).

34. Granados-Fuentes, D., Lambert, P., Simon, T., Mennerick, S. & Herzog, E.D. GABA. Proc Natl Acad Sci U S A 121, e2400339121 (2024).

35. Myung, J., et al. GABA-mediated repulsive coupling between circadian clock neurons in the SCN encodes seasonal time. Proc Natl Acad Sci U S A 112, E3920–3929 (2015).

36. Rohr, K.E., et al. Seasonal plasticity in GABA. Elife 8(2019).

37. Liu, C. & Reppert, S.M. GABA synchronizes clock cells within the suprachiasmatic circadian clock. Neuron 25, 123–128 (2000).

38. Ralph, M.R. & Menaker, M. Bicuculline blocks circadian phase delays but not advances. Brain Res 325, 362–365 (1985).

39. Ralph, M.R. & Menaker, M. GABA regulation of circadian responses to light. I. Involvement of GABAA-benzodiazepine and GABAB receptors. J Neurosci 9, 2858–2865 (1989).

40. Albus, H., Vansteensel, M.J., Michel, S., Block, G.D. & Meijer, J.H. A GABAergic mechanism is necessary for coupling dissociable ventral and dorsal regional oscillators within the circadian clock. Curr Biol 15, 886–893 (2005).

41. Huang, Z.Z., et al. mir-500-Mediated GAD67 Downregulation Contributes to Neuropathic Pain. J Neurosci 36, 6321–6331 (2016).

42. Leppek, K., Das, R. & Barna, M. Functional 5’ UTR mRNA structures in eukaryotic translation regulation and how to find them. Nat Rev Mol Cell Biol 19, 158–174 (2018).

43. Cao, R., et al. Light-regulated translational control of circadian behavior by eIF4E phosphorylation. Nat Neurosci 18, 855–862 (2015).

44. Doi, M., et al. Non-coding cis-element of Period2 is essential for maintaining organismal circadian behaviour and body temperature rhythmicity. Nat Commun 10, 2563 (2019).

45. Takahashi, J.S. Transcriptional architecture of the mammalian circadian clock. Nat Rev Genet 18, 164–179 (2017).

46. McManus, D., et al. Cryptochrome 1 as a state variable of the circadian clockwork of the suprachiasmatic nucleus: Evidence from translational switching. Proc Natl Acad Sci U S A 119, e2203563119 (2022).

47. Morf, J., et al. Cold-inducible RNA-binding protein modulates circadian gene expression posttranscriptionally. Science 338, 379–383 (2012).

48. Park, J.W., et al. Circulating blood eNAMPT drives the circadian rhythms in locomotor activity and energy expenditure. Nat Commun 14, 1994 (2023).

49. Chiang, C.K., et al. The proteomic landscape of the suprachiasmatic nucleus clock reveals large-scale coordination of key biological processes. PLoS Genet 10, e1004695 (2014).

50. Deery, M.J., et al. Proteomic analysis reveals the role of synaptic vesicle cycling in sustaining the suprachiasmatic circadian clock. Curr Biol 19, 2031–2036 (2009).

51. Mito, M., Mishima, Y. & Iwasaki, S. Protocol for Disome Profiling to Survey Ribosome Collision in Humans and Zebrafish. STAR Protoc 1, 100168 (2020).

52. Miyake, T., et al. Minimal upstream open reading frame of Per2 mediates phase fitness of the circadian clock to day/night physiological body temperature rhythm. Cell Rep 42, 112157 (2023).

53. Sasaki, L., et al. Intracrine activity involving NAD-dependent circadian steroidogenic activity governs age-associated meibomian gland dysfunction. Nat Aging 2, 105–114 (2022).

54. Ingolia, N.T., et al. Ribosome profiling reveals pervasive translation outside of annotated protein-coding genes. Cell Rep 8, 1365–1379 (2014).

55. Huang, d.W., Sherman, B.T. & Lempicki, R.A. Systematic and integrative analysis of large gene lists using DAVID bioinformatics resources. Nat Protoc 4, 44–57 (2009).

56. Chen, E.Y., et al. Enrichr: interactive and collaborative HTML5 gene list enrichment analysis tool. BMC Bioinformatics 14, 128 (2013).

57. Kuleshov, M.V., et al. Enrichr: a comprehensive gene set enrichment analysis web server 2016 update. Nucleic Acids Res 44, W90–97 (2016).

58. Doi, M., et al. Gpr176 is a Gz-linked orphan G-protein-coupled receptor that sets the pace of circadian behaviour. Nat Commun 7, 10583 (2016).

